# Identification of Pex34p as a component of the peroxisomal de novo biogenesis machinery in yeast

**DOI:** 10.1101/2021.05.31.446392

**Authors:** Juliane Radke, Shirisha Nagotu, Wolfgang Girzalsky, Anirban Chakraborty, Markus Deckers, Maya Schuldiner, Einat Zalckvar, Ralf Erdmann

## Abstract

Cells can regulate the abundance and composition of peroxisomes to adapt to environmental changes. In the baker’s yeast, *S. cerevisiae*, peroxisomes represent the only site for degradation of fatty acids. Hence, it is not surprising that growth of yeast cells on oleic acid results in a massive proliferation of peroxisomes. New peroxisomes can form either by division of pre-existing peroxisomes or de novo in a Pex25p-dependent process with the involvement of the Endoplasmic Reticulum (ER). In search for further factors involved in de novo formation of peroxisomes, we screened ~6,000 yeast mutants that were depleted of peroxisomes by conditional inhibition of *PEX19* expression. Screening the mutants for the reappearance of peroxisomes upon expression of *PEX19* identified Pex34p, in addition to the well-known component Pex25p, as crucial determinants for de novo biogenesis. Pex34p interacts with Pex19p and with different Peroxisomal Membrane Proteins (PMPs) in a *PEX19*-dependent manner. Depletion of Pex34p results in reduced numbers of import-competent peroxisomes formed de novo and Pex3p is partly retained and distributed in ER-like structures. We suggest that Pex25p and Pex34p are both required to maintain peroxisome number in a cell and that they perform non-redundant roles in the de novo formation of peroxisomes.

---

Peroxisomes are widely distributed among eukaryotic organisms. They participate in a variety of metabolic processes, which are related to the metabolism of lipids and reactive oxygen species (Walter & Erdmann, 2019). The broad range of functions underlies the importance of peroxisomes for human health, as indicated by the plethora of peroxisomal disorders as well as the involvement of peroxisomes in neurodegenerative diseases, obesity, cancer, and age-related disorders (Islinger et al., 2018). Peroxisome abundance is strictly regulated by the rates of organelle formation, division and turnover (Nordgren et al., Wróblewska & van der Klei, 2019). While peroxisomes are degraded by only one main route, pexophagy (Eberhart & Kovacs, 2018, Nordgren et al.), they can be formed by two mechanisms: either by growth and division of pre-existing organelles (fission) or by de novo budding from the ER (Hettema et al., 2014).

Peroxisome fission shares some components with mitochondrial and vacuolar fission. The dynamin-like proteins Vps1p and Dnm1p as well as the Dnm1p-interacting proteins Fis1p, Mdv1p and Caf4p have all been shown to be required for peroxisome fission in *S. cerevisiae* (Hoepfner et al., 2001, Kuravi et al., 2006, Motley et al., 2008). In addition, a role for the Pex11p family of proteins in division has been established (Erdmann & Blobel, 1995, Rottensteiner et al., 2003, Tam et al., 2003). In *S. cerevisiae*, deletion or overexpression of these proteins leads to a reduced or enhanced number of peroxisomes, respectively (Rottensteiner et al., 2003, Vizeacoumar et al., 2003). A conserved role for Pex11p in peroxisome fission has been established in yeast, plants, and mammalian cells. Pex25p and Pex27p belong to the Pex11p family in *S. cerevisiae* (Rottensteiner et al., 2003, Smith et al., 2002, Tam & Rachubinski, 2002). Recent studies, however, point towards several different roles for these proteins in peroxisome biogenesis. Pex11p was shown to be required for peroxisome membrane elongation in *Hansenula polymorpha* (Opalinski et al., 2011) and also for re-organization of peroxisome membrane proteins (Cepińska et al., 2011). However, Pex25p plays a role in the *de novo* formation of peroxisomes as shown in *H. polymorpha* (Saraya et al., 2011) and *S. cerevisiae* (Huber et al., 2012).

Pex34p has been identified as an interaction-partner of the Pex11p-family proteins and is required to maintain peroxisome number in *S. cerevisiae* (Tower et al., 2011). Moreover, Pex34p was also identified as a contact site tether between peroxisomes and mitochondria (Shai et al., 2018). Along this line, overexpression of Pex34p suppresses impaired acetate utilization in yeast lacking the mitochondrial aspartate/glutamate carrier Agc1p (Chalermwat et al., 2019). The formation of peroxisome-mitochondria contact sites from *PEX34* overexpression may facilitate function of an alternative redox shuttle through direct movement of citrate from peroxisomes to mitochondria (Chalermwat et al., 2019).

A major contribution to our understanding of peroxisome biogenesis was the genetic identification of the thirty-seven known peroxisome biogenesis factors (peroxins) (Erdmann & Kunau, 1992). Two central peroxisome biogenesis factors in yeast are Pex3p and Pex19p that together with Pex16p in higher eukaryotes seem to be directly involved in the biogenesis of the peroxisomal membrane(Jansen & van der Klei, 2019, Schrader & Pellegrini, 2017). Accordingly, mutants in Pex3p, Pex19p or Pex16p lack typical morphologically detectable peroxisomal structures (Islinger et al., 2018, Yuan et al., 2016). Peroxisomes reappear after genetic complementation as demonstrated for mammalian cells with a defect in PEX3, PEX16 or PEX19 or yeast mutants of *PEX3* and *PEX19*, indicating that they play an important role not only for peroxisomal targeting of membrane proteins but also for *de novo* formation of peroxisomes (Hettema & Motley, 2009). Pex19p, originally identified as a prenylated protein (PxF) (James et al., 1994, Kammerer et al., 1997) or a housekeeping gene product (HK33) (Braun et al., 1994), displays a high binding affinity for a variety of peroxisomal membrane proteins (PMP) (Jansen & van der Klei, 2019). This gave rise to the idea that Pex19p acts as an import receptor and/or chaperone that binds PMPs in the cytosol, prevents their aggregation and degradation and delivers them to the peroxisomal membrane where it docks to Pex3p (Fang et al., 2004, Hettema et al., 2000, Jones et al., 2004, Shibata et al., 2004). Moreover, Pex19p-function is linked to an early step in peroxisome biogenesis namely the transfer of Pex3p from the ER to peroxisomes (Chen et al., 2014, Matsuzono & Fujiki, 2006). It is likely that additional proteins are involved in this process but have not yet been identified.

Here, we aimed to identify components required for peroxisome *de novo* biogenesis. We performed a genome-wide screen of mutants that delayed the appearance of peroxisomes after being depleted of peroxisomes and then allowed to reform them. We found that in addition to the known biogenesis factor Pex25p and other novel components also Pex34p came up as a crucial determinant. Our data show that Pex25p and Pex34p perform multiple functions in peroxisome biogenesis and maintenance, including non-redundant roles in the *de novo* formation of peroxisomes.

## Materials and Methods

### Strains and growth conditions

The *S. cerevisiae* strain UTL-7A (*MATa ura3-52, trp1, leu2-3/112*) was used as wild-type strain for the generation of several isogenic deletion strains as described previously (Güldener et al., 1996). Yeast strains used are listed in Table 1. Yeast complete (YPD) and minimal media (SD) used have been described previously (Erdmann et al., 1989). YNO medium contained 0.1% oleic acid, 0.05% Tween 40, 0.1% yeast extract and 0.67% yeast nitrogen base without amino acids, adjusted to pH 6.0. When necessary, auxotrophic requirements were added according to (Ausubel et al., 1992). SD medium with 0.5% galactose instead of glucose was used for induction experiments. The screening procedure was carried out with SD-medium containing 2% galactose.

**Table I.**
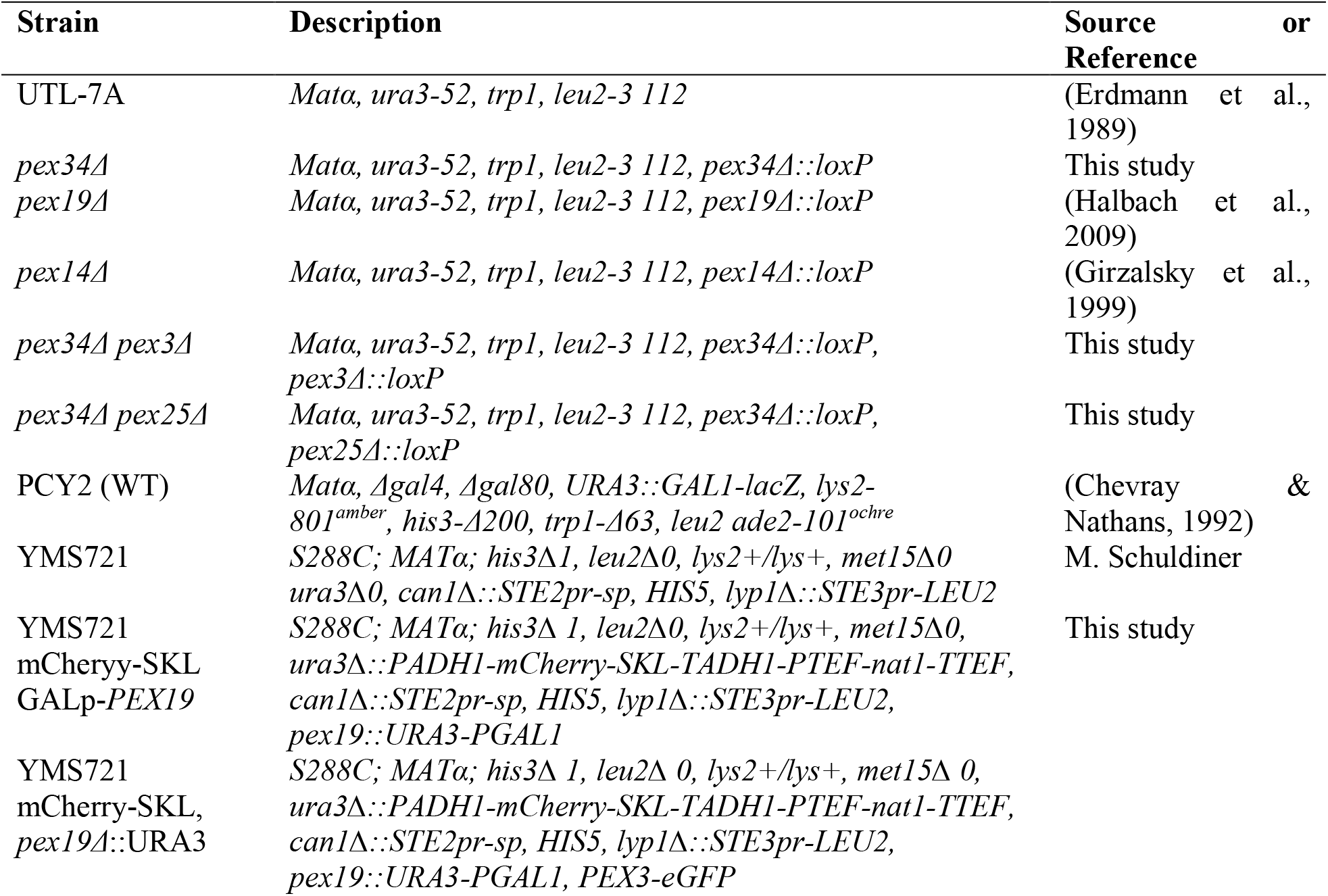
*S. cerevisiae* strains used in this study.

### PEX34

deletion was constructed by homologous recombination based exchange of the open reading frame with KanMX4 (Güldener et al., 1996). LoxP-flanked Kan-MX4 cassettes were obtained by PCR from pUG6. Geneticin resistant colonies were selected and the loxP-KanMX4-loxP was removed by the expression of the Cre-recombinase. Double deletions *pex25*Δ*pex34*Δ and *pex3*Δ*pex34*Δ were made by deleting the respective gene in a *pex34*Δ strain by the exchange of the open reading frame by KanMX4. Primers RE575/862, KU365/KU700 and RE2211/22 were used respectively for *PEX3, PEX19* and *PEX25* deletion cassettes.

The query strain for the screening was generated as follows. First, mCherry-SKL under control of the alcohol-dehydrogenase promotor ADH1 was amplified by PCR from plasmid pAG25 using primers RE4567/RE4568. The obtained PCR product was introduced into the URA3-locus of wildtype YMS721 and isogenic *pex19*Δ strain by homologous recombination. The generated YMS721 mCherry-SKL strain was further used to integrate the *GAL1*-promotor in front of the *PEX19* open reading frame. To this end, the URA-marker as well as the *GAL1*-promotor was amplified by PCR from plasmid pGal-PEX3-GFP with primers RE5292/RE5293. The obtained PCR product was introduced between the native *PEX19-promotor* and the *PEX19* open reading frame by homologous recombination.

### Generation of a screening library

The generated query strain was crossed into the yeast deletion, hypomorphic allele and mini-peroxi collections (Breslow et al., 2008, Gabay-Maskit et al., 2020, Giaever et al., 2002) by the synthetic genetic array method (Cohen & Schuldiner, 2011, Tong & Boone, 2006). This procedure resulted in a collection of haploid strains used for screening mutants that display peroxisomal abnormalities after re-expression of Pex19p.

### Automated high-throughput fluorescence microscopy

The screening collection, containing a total of 6321 strains, were grown for several generations on solid glucose medium to deplete peroxisomes, then for 12 h in 2% galactose medium at 30°C under gently shaking. 5μl of each culture were shifted to 95μl fresh galactose-medium. After additional 4 h growth, cells were visualized during mid-logarithmic growth using an automated microscopy setup as described previously (Breker et al., 2013).

### Plasmid constructions

Plasmids and oligonucleotides used are listed in Table II and III. Plasmid GFP-Pex34 was constructed using the plasmid pUG36 as a template and primers RE1524, RE1582 for PCR. Plasmid pIH024 expressing Pex3-GFP fusion under the control of the Gal4-promotor was constructed as follows: The coding region of Pex3-GFP was amplified by PCR with primers RE1273 and RE1274 and plasmid pIH643 (Heiland & Erdmann, 2005) as template. The PCR product was digested with XhoI/BamHI and cloned into XhoI/BamHI sites of pGal4 (Clontech). For C-terminal GFP-fusion of Pex25, the coding region of Pex25 was amplified from the genome by PCR with primers Pex25-GFP-fw/Pex25-GFP-rev and introduced into EcoRI/SalI sites of pUG35. All constructs were confirmed by DNA sequencing. The GFP-SKL and DsRed-SKL plasmids have been described previously (Brocard et al., 1997, Halbach et al., 2009).

**Table II.** Plasmids used in this study.

| Plasmid | Description | Source / Reference | Primers |
| --- | --- | --- | --- |
| pPC86- | Gal4p(DB) | (Chevray & Nathans, 1992) |  |
| pPC97- | Gal4p(AD) | (Chevray & Nathans, 1992) |  |
| pPC86-PEX19 | Gal4p(DB)-Pex19p | (Rottensteiner et al., 2004) |  |
| pPC86-PEX34 | Gal4p(DB)-Pex34p | This study | RE1524/RE1525 |
| pUG6 | Deletion, Cre-Recombinase | (Güldener et al., 1996) |  |
| pUG36 | GFP | (Niedenthal et al., 1996) |  |
| pUG36-PEX34 | GFP-Pex34p | This study | RE1524/RE1582 |
| pUG35-PEX25 | Pex25p-GFP | This study | Pex25p-GFP-fw/Pex25p-GFP-rev |
| pRS414-Ds-Red-SKL | DsRed-SKL (-trp) | (Halbach et al., 2009) | RE1685/RE1686 |
| pGal-PEX3-GFP | Pex3p-GFP | This study | RE1273/RE1274 |

**Table III.**
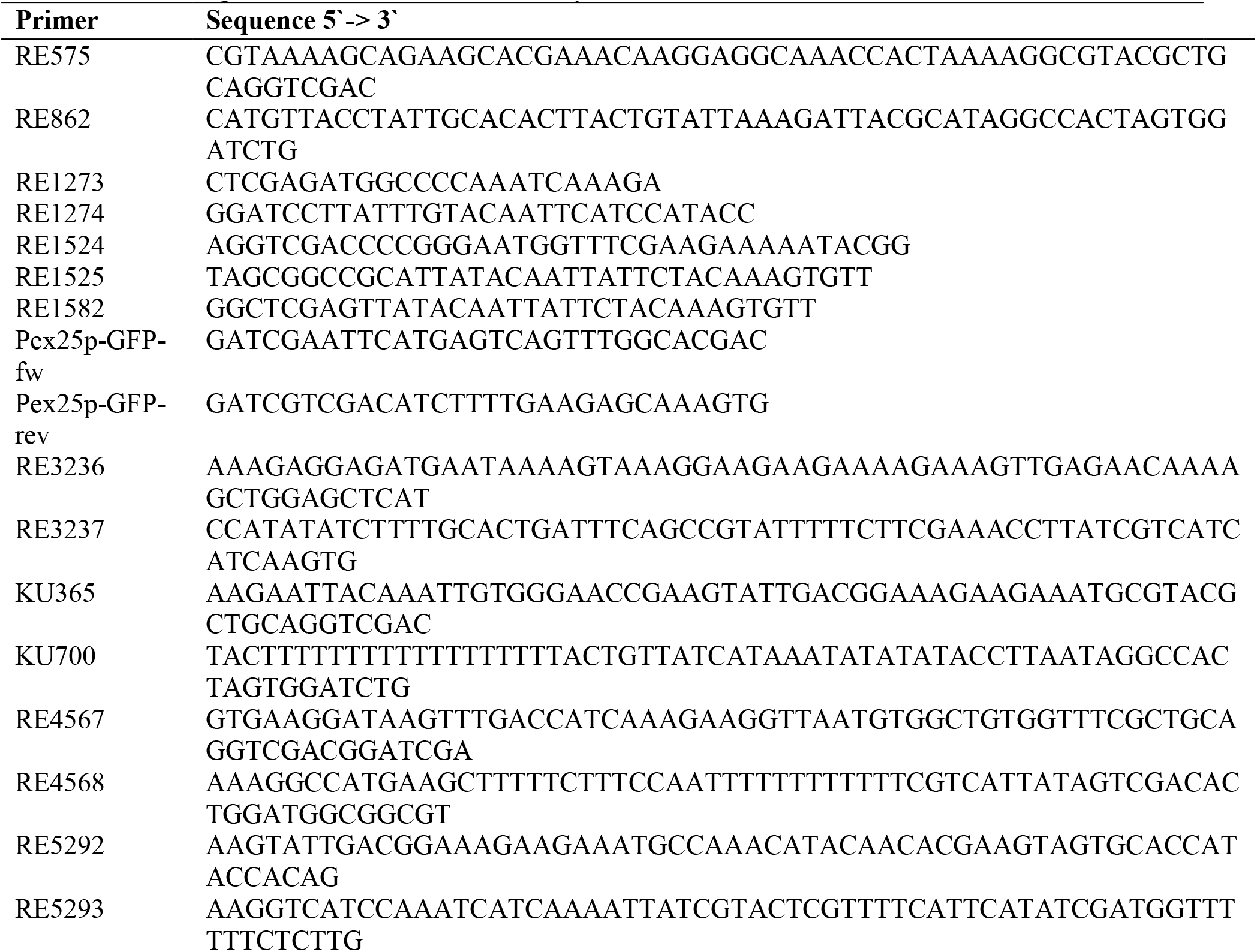
Oligonucleotides used in this study.

### Two-hybrid Analysis

The two hybrid analysis was based on the method of (Fields & Sternglanz, 1994). The tested genes were fused to the DNA-binding domain (Gal-DB) or trans-activating domain (Gal-AD) of Gal4 in the vectors pPC86 and pPC97 (Chevray & Nathans, 1992). Co-transformation of two hybrid vectors into strain PCY2 was performed according to (Gietz & Woods, 1994). Transformed yeast cells were plated on SD synthetic medium without tryptophan and leucine. β-galactosidase filter assays were performed as described earlier (Rehling et al., 1996b).

### Image acquisition

Fluorescence microscopy images were recorded on an AxioPlan 2 microscope (Zeiss, Jena) equipped with a αPlan-FLUAR 100×/1.45 oil objective and an AxioCam MRm camera (Zeiss, Jena) at room temperature. If necessary, contrast was linearly adjusted using the image acquisition software AxioVision 4.8 (Zeiss, Jena).

### Miscellaneous

Immunoblotting was performed with polyclonal rabbit antibodies raised against Pex19p (Götte et al., 1998) and mitochondrial porin (Kerssen et al., 2006).

## Results

### Construction of a conditional Pex19p-mutant strain

Cells deficient in Pex19p are characterized by the lack of peroxisomes, with the consequence that the peroxisomal matrix proteins are mislocalized to the cytosol. New peroxisomes can form *de novo* in these cells, upon reintroduction of *PEX19* (Götte et al., 1998). To carry out a genome wide screen to identify components involved in this process, we first generated a query strain that enables us to control and monitor *de novo* formation of peroxisomes. To this end, the *GAL4*-promotor was integrated in front of the *PEX19*-coding region, replacing the endogenous *PEX19*-promotor (**FIG 1A**) in a yeast strain compatible with automated mating, sporulation and segregant selection. This procedure enabled us to switch off *PEX19-expression* on glucose medium, whereas presence of galactose induced *PEX19* expression. To monitor formation of functional and thus import-competent peroxisomes, the fluorophore mCherry fused to the peroxisomal targeting-signal type 1 (PTS1: SKL) under control of the constitutive ADH1-promotor was additionally integrated into the genome of the query strain (**FIG 1A**).

**Figure 1:**
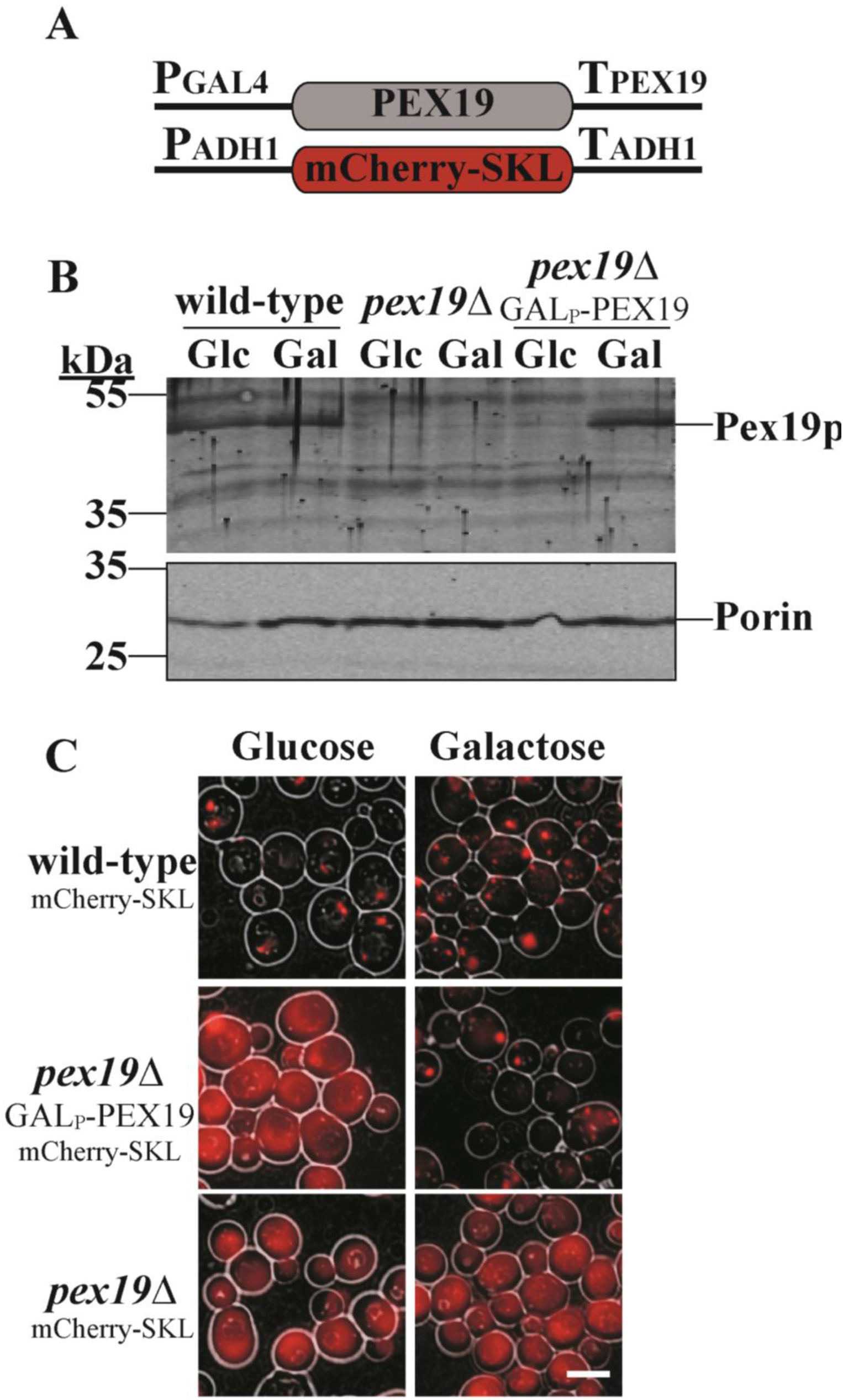
Generation and characterization of the query strain for a genome-wide screen. (A) Schematic representation of the genetic modification of strain YMS721. The GAL4-promotor was introduced by homologous recombination upstream of the *PEX19-*coding region. mCherry fused to PTS1-signal SKL under control of the ADH1-promotor was integrated into the *URA3*-locus. (B) Strains as indicated were first grown to mid log-phase on glucose-medium and then shifted to either glucose (Glc) or galactose (Gal) conditions. After growth for 4 h, whole cell TCA-lysates were prepared. Equal portions were subjected to SDS-PAGE followed by immunoblot analysis using anti-Pex19p and anti-porin (mitochondrial marker protein, served as loading control) antibodies. (C) Indicated cells were grown in glucose medium until mid-log-phase, shifted after a washing step to galactose conditions and grown for 4 h. Prior and after growth on galactose, cells were analyzed by fluorescence microscopy for localization of mCherry-SKL. The scale bar represents 5 μm.

To analyze the regulation of *PEX19*-expression, cells of the control, *pex19*Δ strain and the newly generated query strain (*Gal4-PEX19*) were first grown on glucose-medium, subsequently shifted to either glucose or galactose containing medium and whole cell lysates thereof were analyzed for Pex19p-abundance. Mitochondrial porin served as an internal loading control and did not differ between the analyzed samples. In control cells, the Pex19p-steady-state level was nearly the same under both, glucose- and galactose-conditions, whereas no Pex19p-positive signal was obtained in samples of the *pex19*Δ mutant as expected (**FIG 1B**). In the query strain, Pex19p was not detected in the lysates from glucose-grown cells, indicative of a tight repression. In contrast, the lysates from galactose-induced cells contained similar amounts of Pex19p as wild-type cells. (**FIG 1B**). This clearly demonstrated a tight glucose-repression and efficient galactose-induction of *PEX19* in the query strain.

Although the Pex19p-concentration in our query strain was below the detection level when glucose served as a carbon source, we confirmed that the basal expression is low enough to block peroxisome biogenesis and that growth on glucose-medium for several generations sufficiently depleted peroxisomes of the parental strain. To this end, the query strain was compared with wildtype and *pex19*Δ cells for the presence of import-competent peroxisomes in glucose-grown and galactose-induced cells, monitored by fluorescence microscopy. In line with a peroxisomal localization, a punctuate pattern of the genomically expressed mCherry-SKL was seen in control cells, independent of the growth conditions (**FIG 1C**). As expected for a strain impaired in peroxisome biogenesis (Distel et al., 1996), the synthetic peroxisomal marker was mislocalized to the cytosol and thus caused a diffuse staining in the *pex19*Δ strain. While under glucose conditions the query strain behaved like a *pex19*Δ mutant, red puncta were monitored when *PEX19*-expression was induced by the presence of galactose (**FIG 1C**). We quantified that no more than 6 % of the query cells maintained import-competent peroxisomes even under glucose conditions (and those that did had no more than a single one) making this strain highly suitable to screen for components of the peroxisomal *de novo* biogenesis.

### Screening for components required for the peroxisomal de-novo biogenesis

The created query strain was compatible with the Synthetic Genetic Array (SGA) automated mating approach in yeast (Cohen & Schuldiner, 2011, Tong & Boone, 2006). To this end, this strain was crossed into the nearly 5000 strains of the deletion library (Giaever et al., 2002) and more than 1000 strains of the DAmP (Decreased Abundance by mRNA Perturbation) hypomorphic allele library for essential genes. Following sporulation and selection of haploids, about ~6000 unique haploid strains were obtained, each with a deletion of a single yeast gene and harboring *GAL4-PEX19* as well as mCherry-SKL (**FIG 2**). To perform the screen, the generated screening library was first grown on glucose medium with intermediate medium changes to force peroxisome depletion. Thereafter, cells were shifted to galactose medium and grown for 16 h again with intermediate medium change to induce Pex19p-expression. Finally, cells were subjected to microscopic inspection and automated image acquisition. Using an automated high-content microscopy setup specifically configured for this purpose (Breker et al., 2013), three images of one focal plane were recorded for each mutant covering hundreds of cells per strain. Images were analyzed for the appearance of fluorescence labelling differing from that of control cells. Analyzed strains were grouped into those with no obvious difference to the control and those that contain mature peroxisomes that differed from control cells by number or size. Finally, we found strains that had complete mislocalization of the peroxisomal marker mCherry-SKL to the cytosol or a mixed phenotype (**FIG 2**).

**FIGURE 2:**
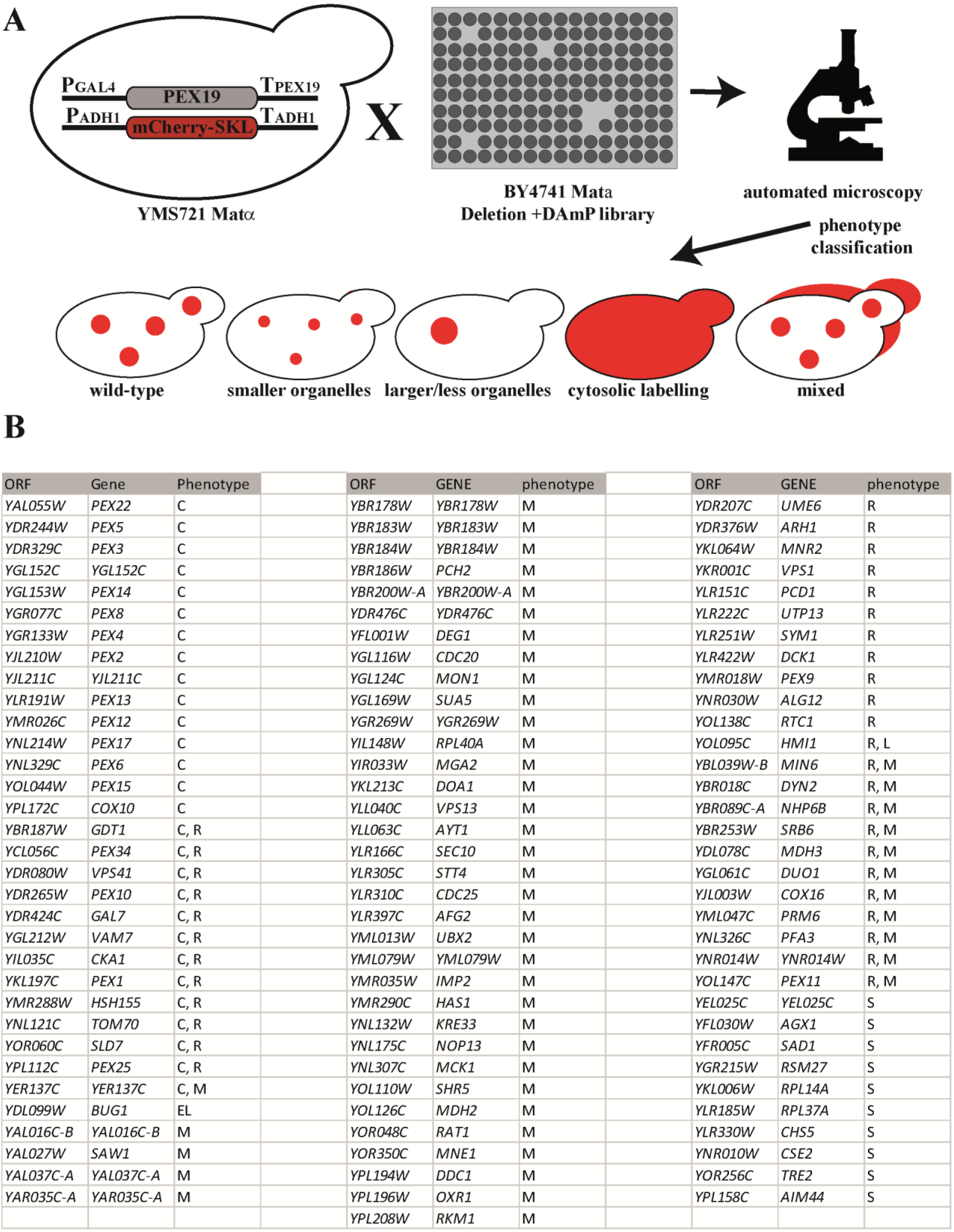
A Genome-wide screen for components involved in peroxisomal de-novo biogenesis. (A) Schematic workflow of the genome-wide screen. GAL4-*PEX179*/mCherrv-SKL were integrated into yeast strain YMS721 to generate the query strain. This was crossed with the indicated deletion/DAmP libraries, sporulated and selected for haploid deletion-strains harboring both the GAL4-*PEX19*/mCherry-SKL as well as single gene-deletions. Strains were analyzed by automated microscopy and were grouped into indicated phenotypes based on the appearance of the peroxisomal marker mCherry-SKL. B) List of mutants that display a localization of mCherry-SKL different from that of wild-type cells. C-cytosolic distribution of the reporter; S – small peroxisomes; L – large peroxisomes; R – reduced number of peroxisomes; El – elongated peroxisomes; M – mixed phenotype

In total, 100 strains were identified, which differed from the control in peroxisome appearance (**Fig 2B; Supplementary table 1**) and were defined as hits. Eighteen of them turned out to be affected in one of the known PEX-genes, which encode peroxisomal assembly factors (Islinger et al., 2010). In two strains *(YGL152C, YJL211C)*, the corresponding gene is considered to be dubious but is overlapping a PEX gene: *PEX14* and *PEX2* respectively, and the deletion would have deleted them as well. It had been demonstrated previously that yeast mutants lacking peroxisomes require the presence of Pex3p and Pex25p to regenerate this organelle *de novo* (Huber et al., 2012, Saleem et al., 2008). Accordingly, *pex3*Δ and *pex25*Δ both displayed eye-catching phenotypes. The *pex3*Δ mutant cells displayed a distinct cytosolic labelling of mCherry-SKL, which is typical for a strain affected in peroxisomal matrix protein import or biogenesis (**FIG 3**; (Distel et al., 1996)). The *pex25*Δ cells showed less peroxisomes and a partial mislocalization of the peroxisomal marker to the cytosol (**FIG 3**). The fact that we identified Pex3p and Pex25p, two known components of the de novo biogenesis (Huber et al., 2012, Saleem et al., 2008), and the finding that the hits of our screen cover almost all components known to be required for import of type1 peroxisomal matrix-proteins or peroxisomal appearance, demonstrated the validity of the screen.

**FIGURE 3:**
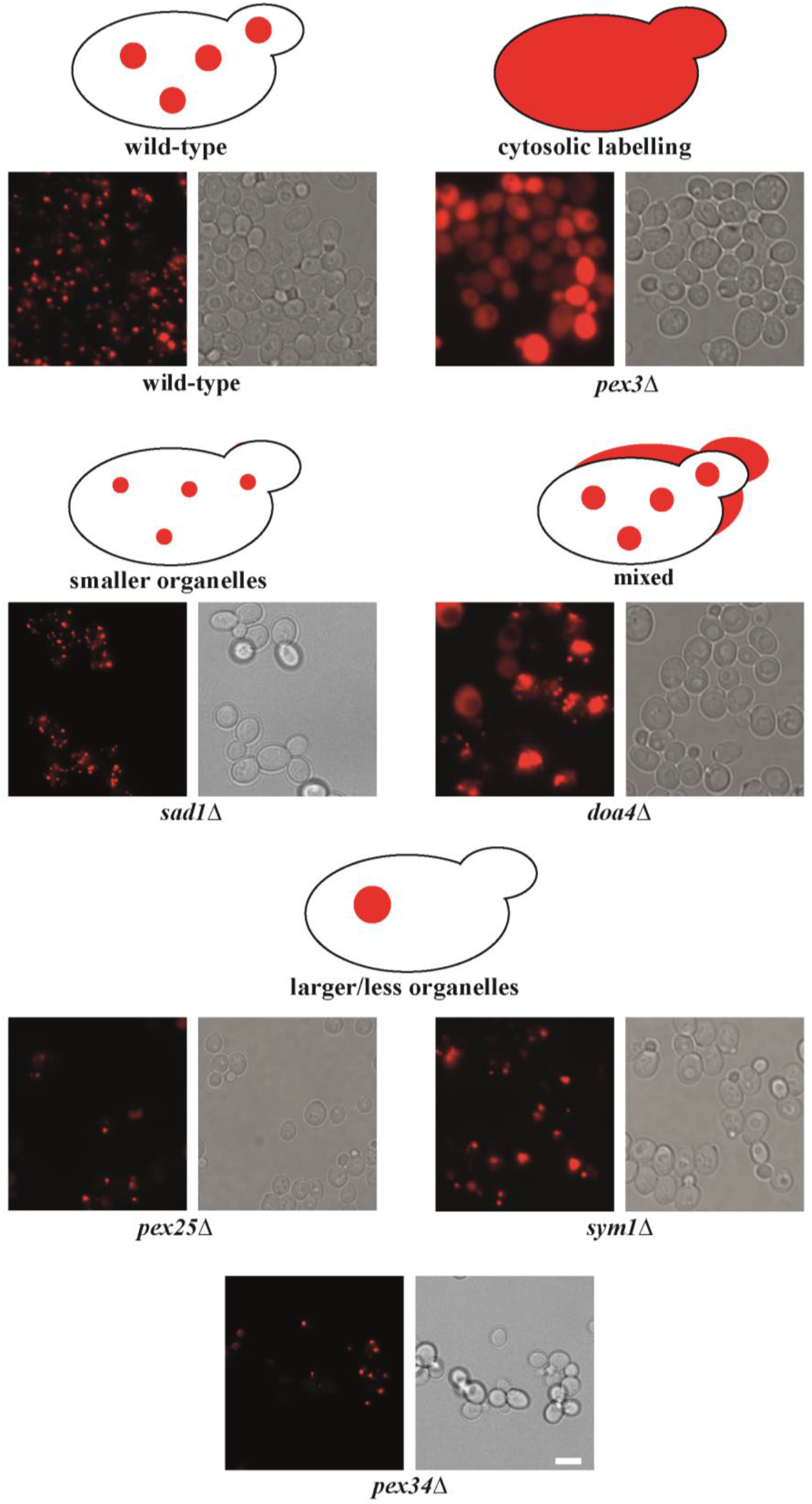
Examples of hits from the genome wide-screen. Examples of mutants from the genome-wide screen. Images of strains from the genome-wide screen containing GAL4-PEX19 and mCherry-SKL. Strains were first grown on glucose-medium and then shifted to galactose-medium to induce Pex19p expression. Images were taken after 16h galactose-induction with intermediate medium changes. The scale bar represents 5 μm.

However, there were many additional hits that were previously not identified as taking part in this process but have strong connections to peroxisomes and may be important to further study. For example: A deletion of *SAD1* caused the appearance of multiple but smaller peroxisomes. Sad1p is involved in pre-mRNA-spicing (Lygerou et al., 1999). In this respect, it is interesting to note that several of the genes identified in the screen contain introns (*NHP6B, PCH2, RPL14A, RPL37A and RPL40A*). A deletion of *DOA4* caused a mixed phenotype of normal import and a partial import defect (**FIG 3**). Doa4p was shown to be involved in cleavage of ubiquitin chains and is required for turnover of the PTS2 co-receptor Pex18p, which also is ubiquitinated (Papa & Hochstrasser, 1993, Purdue & Lazarow, 2001). Growth-rate of *doa4*Δ cells was shown to be reduced on both, glucose and oleic-acid conditions, and matrix protein import is affected (Debelyy et al., 2011).

An additional interesting hit is Mdh2p whose deletion caused a mixed phenotype mixed phenotype of normal import and a partial import defect. Mdh2 was described as an enzyme involved in both, the glyoxylate cycle and fatty acid β-oxidation (Kunze et al., 2006). Initially described as a cytosolic protein, recent data demonstrated a fraction of Mdh2p to be localized to peroxisome, depending on Pex5p as well as Mdh3p (Gabay-Maskit et al., 2020). Deletion of *<u>YAL016C-B</u>* - a dubious open reading frame that overlaps with the promotor region of *TPD3* caused a mixed phenotype similar to those observed in *doa4*Δ and *mdh2*Δ cells. Tpd3p represents a subunit of the PP2A complex, which is a subgroup of the PPP-(serine/threonine-specific phosphoprotein phosphatases) family. These phosphatases are conserved among eukaryotes and a peroxisomal localization was reported for the protein from Arabidopsis thaliana (Kataya et al., 2019, Lillo et al., 2014). Recently, regulation of peroxisomal matrix protein import via phosphorylation of Pex14p, a key component of the import machinery, was reported, is may be that yeast Tpd3p in involved in this process (Schummer et al., 2020). Shr5p (Erf4) is involved in the palmitoylation and subcellular localization of yeast Ras protein (Zhao et al., 2002). *shr5*Δ appeared in our screen with a mixed phenotype. As Pex19p itself is farnesylated, the observed phenotype might be due to less or lacking Pex19p-modification. We also identified *vps1*Δ by a reduced peroxisome number, a phenotype, which was also reported previously (Hoepfner et al., 2001, Kuravi et al., 2006). Vps1p is required for fission of existing peroxisomes and acts together with Mdm1p (Motley & Hettema, 2007). An additional interesting hit of our screen with a mixed phenotype was *VPS13*. In mammals, the VPS13 gene family consists of VPS13A-D. Recently, VPS13D was shown to be involved in regulation of peroxisomal biogenesis and bridging of the ER to mitochondria and peroxisomes (Baldwin et al., 2021, Guillén-Samander et al., 2021). Deletion of *sym1*Δ gave a dramatic phenotype of reduced number of peroxisomes (**FIG 3**). Sym1p belongs to the PXMP2 protein-family, named by human PXMP2, a channel forming protein of mammalian peroxisomes (Rokka et al., 2009). Recently, Pex37p of *Hansenula polymorpha* was identified, which also belongs to this protein-family. A deletion of *PEX37* displays, like *sym1*Δ, a reduced number of peroxisomes (Singh et al., 2019), indicating that Sym1p might be the functional orthologue of Pex37p in *S. cerevisiae*.

For a more detailed analysis, we chose to focus on one of the most severe hits, the gene *PEX34*. Pex34p was originally reported to function in concert with the Pex11p-protein family members Pex11p, Pex25p, and Pex27p to control the peroxisome populations of cells under conditions of both peroxisome proliferation and constitutive peroxisome division (Rottensteiner et al., 2003, Tam et al., 2003, Tower et al., 2011). However, Pex25p also plays a role in the de novo synthesis of peroxisomes (Huber et al., 2012). As Pex34p has been reported to interact with Pex25p (Huber et al., 2012), we considered that both proteins might function together in peroxisomal de novo synthesis.

### Interaction of Pex34p-Pex19p

Pex34p was originally identified as a peroxisomal membrane protein, which are typically bound by the import receptor Pex19p prior to peroxisomal targeting (Rottensteiner et al., 2004, Tower et al., 2011). The targeting signal for peroxisomal membrane proteins is defined as a Pex19p-binding motif in conjunction with a transmembrane segment or a peroxisomal anchor sequence (Girzalsky et al., 2006, Rottensteiner et al., 2004). The use of topology prediction programs revealed that Pex34p contains two transmembrane segments ((Tower et al., 2011), **FIG 4A**). Moreover, we identified two regions (amino acids 58-72 and 133-143) fitting to the consensus sequence of putative Pex19p-binding sites, which are located in close proximity to the *in silico* predicted transmembrane spanning regions (**FIG 4A**). Thus, Pex34p seems to be a typical Pex19p-cargo. To support this assumption, we investigated whether Pex19p directly interacts with Pex34p by means of a yeast two-hybrid system. Gal4p-fusions of activation domain (AD)-Pex34p and DNA-binding domain (DB) of Pex19p were co-expressed in the *S. cerevisiae* Y2H-host strain PCY2 and interactions were monitored by β-galactosidase filter assays. In line with the idea of Pex34p being a cargo of Pex19p, a clear interaction was indicated by our Y2H-result (**FIG 4B**).

**Figure 4:**
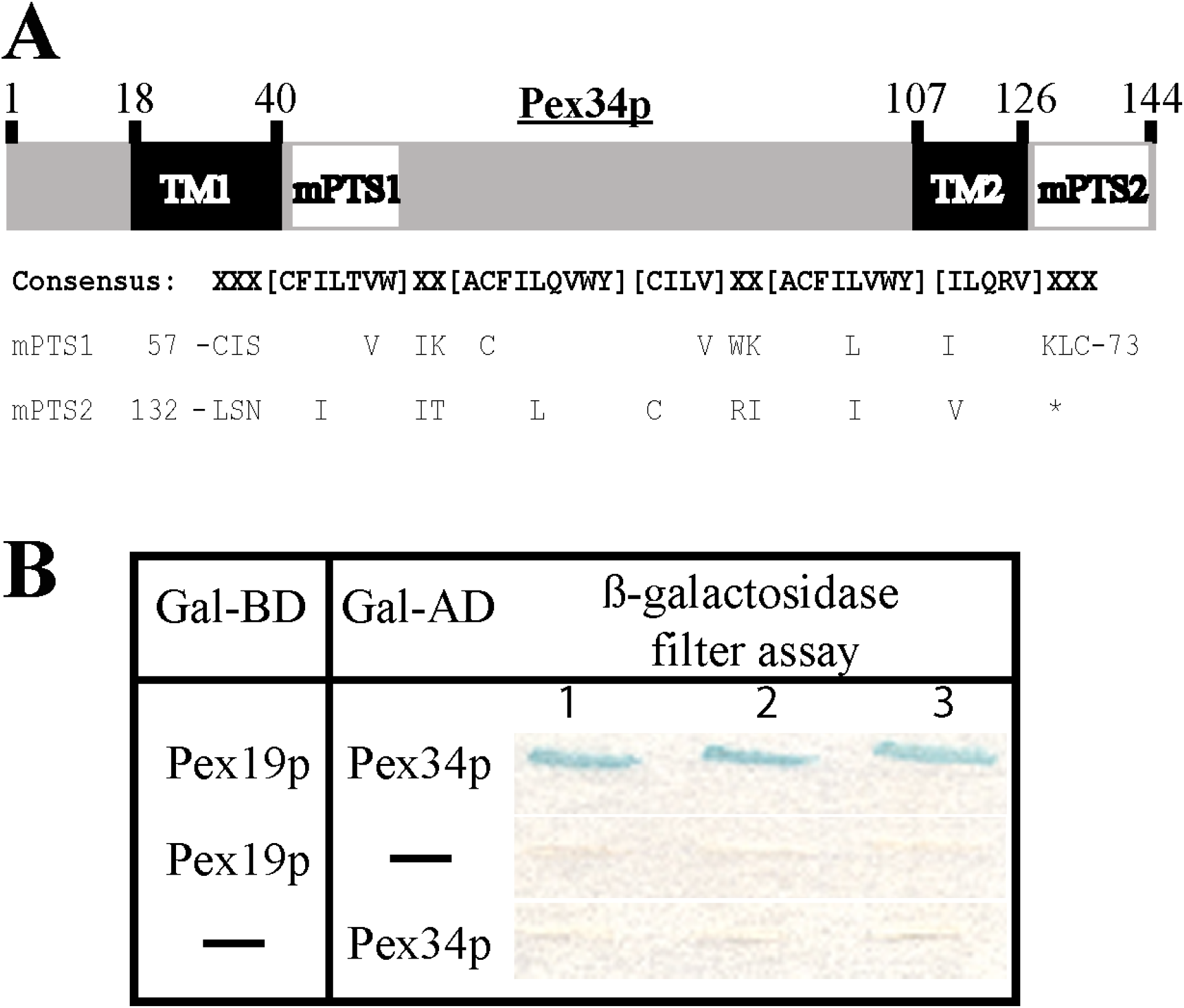
Pex34p interacts with Pex19p. (A) Schematic representation (upper panel) of Pex34p with numbers representing position of corresponding amino-acid residues. Hypothetic transmembrane segments (TM1 and TM2) and Pex19p-binding motives (mPTS1 und mPTS2) are highlighted. Lower panel: Alignment of the consensus sequence for Pex19p-binding motif (Rottensteiner et al., 2004) with the two identified motifs (mPTS1 and mPTS2) within Pex34p. Position are indicated by numbers, the asterisk indicates the stop-codon. (B) Double transformants of the two-hybrid reporter strain PCY2 expressing the indicated GAL-fusion proteins, were selected and β-galactosidase activity was determined by a filter assay with X-Gal as substrate. Three representative independent double transformants are shown.

### Pex34p is required for proliferation of newly formed peroxisomes in pex3Δpex34Δ cells

Considering a function of Pex34p in peroxisomal *de novo* synthesis, we had a closer look in re-appearance of peroxisomes in selected single and double mutant strains. To this end, we applied an established assay for *de novo* synthesis based on a plasmid-encoded *PEX3* under regulation of the *GAL4*-promotor (Hoepfner et al., 2005). Similarly to Pex19p, the role of Pex3p in *de novo* biogenesis of peroxisomes has been established (Aranovich et al., 2014, Ma et al., 2011, van der Zand et al., 2010). This assay also allowed the direct monitoring of the expression and localization of GFP-tagged Pex3p. To this end, a *pex34*Δ*pex3*Δ double deletion was constructed that lacks peroxisomal structures (data not shown). For the *de novo* restoration of peroxisomes in these cells, Pex3p-GFP fusion protein, expressed under the control of the GAL promoter, was reintroduced into the cells. To this end, cells were first pre-cultured on glucose medium, which represses the GAL promoter. Cells were then shifted to medium containing 0.5% galactose for the induction of Pex3p-GFP and initiation of *de novo* peroxisome formation, which was monitored by fluorescence microscopy. The *pex3*Δ cells with GAL-induced Pex3p-GFP served as control. At T0 (just after shifting to galactose containing medium), a detectable Pex3p-GFP signal was not observed in the cells (**FIG 5A**). After 2hrs of induction, Pex3p-GFP puncta were visible, which increased in size and number with time in the *pex3*Δ control cells (**FIG 5A**). In contrast, peroxisome number was significantly reduced in *pex34*Δ*pex3*Δ cells upon reintroduction of Pex3p-GFP (**FIG 5A, B**). Pex3p-GFP was observed as a single spot after two hours of induction and spot number did not significantly increase with induction-time (**FIG 5A, B**). The single punctate structure that contained Pex3p-GFP was confirmed to be an import competent peroxisome as shown by co-labelling with DsRed-SKL (**FIG 5C**). Important to note, we also observed a fraction of Pex3-GFP in a reticulated membrane structure, presumably the ER (**FIG 5A**). We take this finding as indication of a partial targeting to or longer retention of Pex3p in the ER.

**Figure 5:**
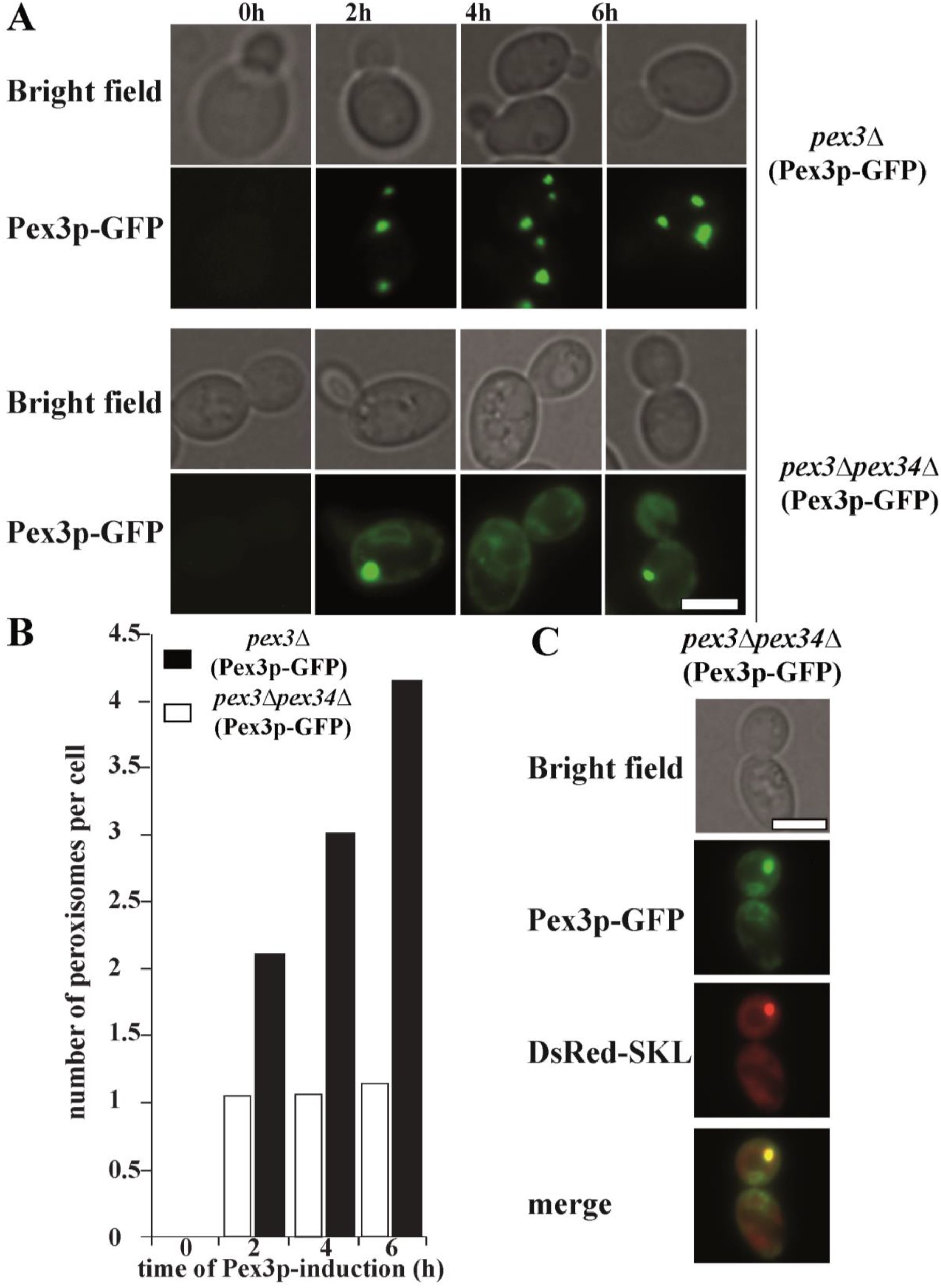
Pex34p is required for efficient re-introduction of peroxisomes. (A) Pex3p-GFP plasmid under the control of the inducible GAL4-promoter was transformed into *pex3Δ* and *pex3Δpex34*Δ cells. Cells were precultured on repressing conditions (glucose medium) and then shifted to medium with 0.5% galactose to induce Pex3p-GFP and monitor peroxisome formation at indicated time points by fluorescence microscopy. (B) For quantification, peroxisomes of hundred cells of each strain (depicted in A) at indicated time-points were counted for two different experiments and the average number is displayed. In *pex3Δ* cells, after two hours of cultivation on galactose medium, Pex3p-GFP was observed in punctuate peroxisomal structures, which proliferated upon prolonged cultivation. In *pex3Δpex34Δ* cells after two hours of induction, Pex3p-GFP localized to a single puncta and additionally to ER-like structures. (C) The single Pex3p-GFP puncta present in *pex3Δpex34Δ* cells upon Pex3p-GFP expression co-localized with the peroxisomal matrix marker DsRed-SKL and thus represent import-competent peroxisomes. The scale bar in panels A and C represents 5 μm.

### Reintroduction of Pex34 in pex25Δpex34Δ cells does not fully restore peroxisome formation

Upon induction, mutants lacking either Pex25p or Pex34p are characterized by a reduced number of import-competent peroxisomes (Tower et al., 2011, Vizeacoumar et al., 2003). In contrast, the phenotype of the corresponding double mutant is more severe with most cells being characterized by a greater reduction in peroxisome number when compared to the single deletion strains and a mislocalization of at least one peroxisomal matrix protein to the cytosol (Tower et al., 2011). Moreover, Pex25p has been reported to be required for the *de novo*-formation of peroxisomes (Huber et al., 2012, Saraya et al., 2011). To gain more insight into a putative concerted function of Pex34p and Pex25p, we first compared the phenotype of *pex34*Δ*pex25*Δ cells upon inducing and non-inducing conditions. Upon induction of peroxisome proliferation by growth on oleic-acid medium, a mixed phenotype was observed. As judged by the localization of the peroxisomal marker GFP-SKL, this marker is mislocalized to the cytosol in a population of oleic acid-induced *pex34*Δ*pex25*Δ cells, indicating that the double mutant exhibits an import defect for peroxisomal matrix proteins (**FIG 6A**). However, this phenotype is not evident on glucose medium, where most of the cells have labeled peroxisomes (**FIG 6A**). The less pronounced phenotype of the *pex34*Δ*pex25*Δ mutant grown under non-inducing conditions might indicate that a low rate of peroxisome formation can still take place normally in the absence Pex34p and Pex25p. On the other hand, the import defect that is seen under inducing conditions might indicate that a higher rate of peroxisome formation requires the presence of both proteins.

**Figure 6:**
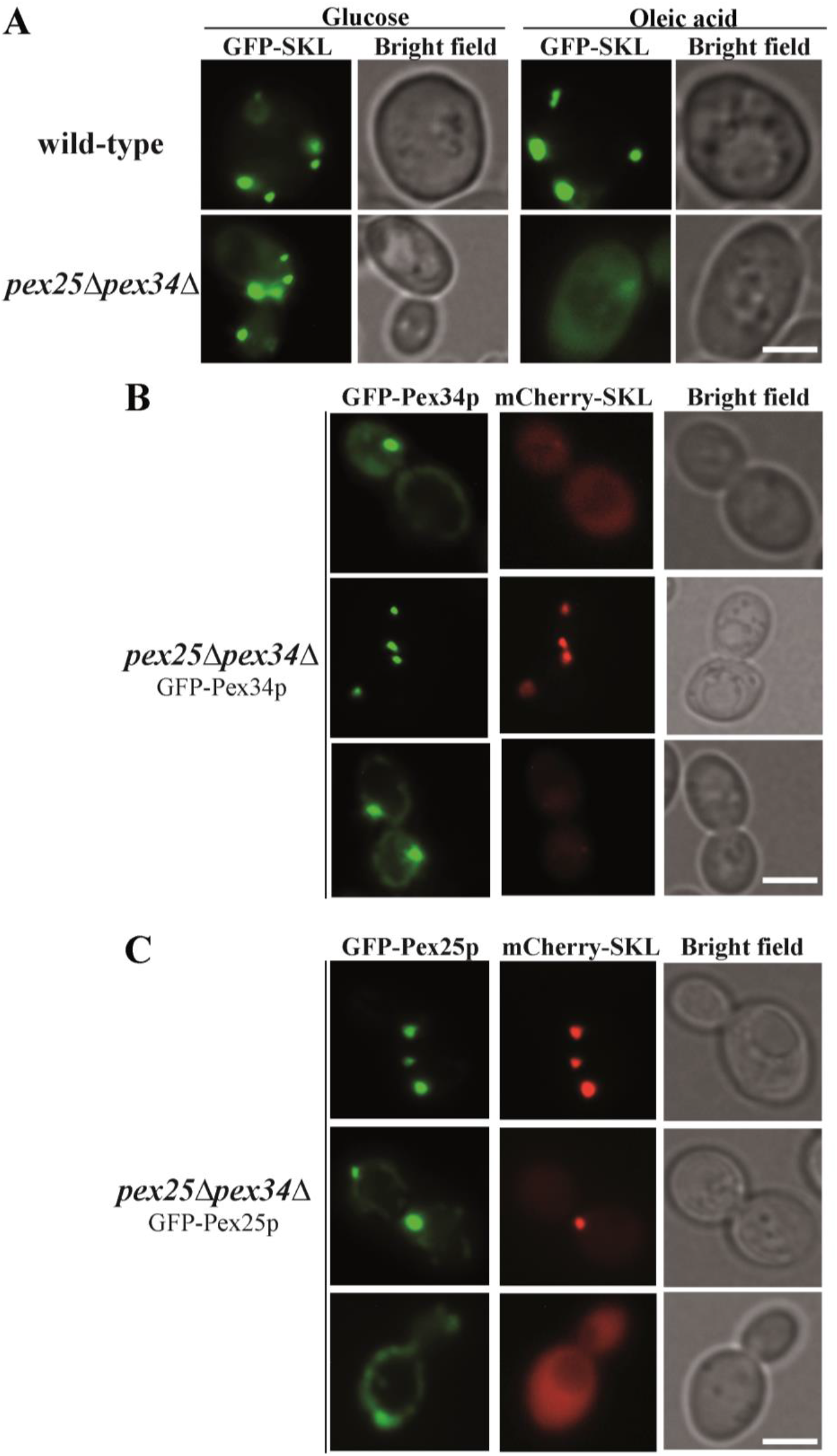
Reintroduction of GFP-Pex34p or Pex25p-GFP in *pex34*Δ*pex25*Δ cells does not restore peroxisome formation. (A) GFP-SKL plasmid was transformed to wild-type and *pex34*Δ*pex25*Δ cells. Peroxisome morphology and abundance of cells grown on glucose and on oleic acid was analyzed by fluorescence microscopy. Cells were precultured on glucose and were shifted to glucose or oleic acid medium. Many *pex34*Δ*pex25*Δ cells showed mislocalization of GFP-SKL when grown on oleic acid medium. The scale bar represents 2.5 μm. (B) Plasmid GFP-Pex34p or (C) Pex25p-GFP was transformed to *pex34*Δ*pex25*Δ cells and GFP-localization was analyzed by fluorescence microscopy. Cells that displayed a normal peroxisomal localization of both GFP-Pex34p and Pex25p-GFP also exhibited a normal import of synthetic marker protein mCherry-SKL, indicated by the clear co-localization. In contrast, when Pex25p or Pex34p appeared to be mislocalized, the peroxisomal matrix protein import was affected and the marker proteins accumulated in the cytosol. The scale bar represents 5 μm.

As the phenotypes of the single and double mutants differ significantly, we analyzed their complementation with either of the two proteins. To this end, we reintroduced GFP-Pex34p into *pex34*Δ*pex25*Δ strain to investigate whether the double mutant cells that are obviously devoid of peroxisomes form functional peroxisomes upon complementation of one of the defects. Surprisingly, we again observed a mixed phenotype. In cells containing punctuate mCherry-SKL, co-localization of GFP-Pex34p was observed indicating a peroxisomal localization of the protein (**FIG 6B**). However, in cells with no detectable peroxisomal structures (cytosolic mCherry-SKL), GFP-Pex34p was observed to be localized to single spots. This result indicates that complementation of *pex34*Δ*pex25*Δ cells with Pex34p does not fully rescue the phenotype.

Similarly, we also reintroduced Pex25p-GFP in *pex34*Δ*pex25*Δ cells and analyzed whether the mutant phenotype of these cells could be rescued by Pex25p. However, a similar localization pattern as for Pex34p was observed.

Thus, single deletions of Pex25p or Pex34p results in a reduced number of functional peroxisomes but upon the concurrent depletion of both proteins a fraction of cells is devoid of peroxisomes, a phenotype which cannot be rescued by complementation with either Pex34p or Pex25p. Accordingly, both proteins play non-redundant roles in the de novo formation of peroxisomes.

## Discussion

Peroxisomes are formed by either growth and division of existing organelles or *de novo* with the involvement of the ER and the peroxisomal membrane protein Pex3p (Haan et al., 2006, Hoepfner et al., 2005, Kragt et al., 2005, Titorenko et al., 1997). In yeast cells, growth and division is the major source for new peroxisomes under normal conditions (Motley & Hettema, 2007), whereas under conditions in which peroxisomes are absent (e.g. segregation defect), newly synthesized Pex3p targets to the ER, concentrates at the membrane into foci and buds of as vesicles which later import matrix proteins and transform into mature peroxisomes (Hoepfner et al., 2005). Here, we performed a genome wide screen for strains affected in peroxisomal *de novo* biogenesis. A conditional strain was generated in which Pex19p-expression was induced by presence of galactose and repressed in glucose-containing medium. Growth in glucose resulted in a complete absence of Pex19p and depletion of peroxisomes from the cells (**FIG 1**). Upon induction of *PEX19* expression, these cells were able to form new peroxisomes *de novo*. Combined with about 6000 single-gene deletions, we identified 100 strains, which display peroxisomes that differ from wild-type organelles in terms of number, size or import capacity. Among the hits, a number of strains lacking proteins known to be required for proper peroxisome biogenesis (PEX-proteins) appeared, demonstrating the validity of the screen (**FIG 2, Supplementary table 1**).

One hit of our screen was the *sym1*Δ mutant. This strain differed from the control by a reduced number of peroxisomes upon induction of peroxisomal *de novo* synthesis (**FIG 3**). Sym1p (for “stress-inducible yeast Mpv17”), is a heat-induced gene product, with a role in ethanol metabolism and tolerance and a homologue to mammalian Mpv17 (Antonenkov et al., 2015, Trott & Morano, 2004). Both, Sym1p and Mpv17p belong to the PXMP2-family, which additionally includes Pxmp2p, Mpv17-like protein (M-LP), and Fksg24p (Mpv17-like 2 protein)) in mammals and Yor292p in yeast (Antonenkov et al., 2015). In contrast to the peroxisomal localization of Pxmp2p, the Mpv17p, MP-L, and Mpv17L2p proteins are mitochondrial (Rokka et al., 2009). The same is true for Sym1p, which is an integral membrane protein of the inner mitochondrial membrane (Trott & Morano, 2004). Recently, Pex37p was identified as an additional family member in the methylotrophic yeast *Hansenula polymorpha* (Singh et al., 2019). The corresponding *pex37*Δ strain was not affected in peroxisome biogenesis or proliferation under peroxisome inducing condition. However, loss of Pex37p caused a significant reduction in peroxisome number under peroxisome repressing conditions, like growth on glucose (Singh et al., 2019). The fact that also peroxisome segregation was affected in *pex37*Δ led to the suggestion that in addition to Pex11p and Dnm1p, Pex37p plays a role in peroxisome fission under glucose condition. Except for the fact that Sym1p was shown to be a mitochondrial protein, it plays a role for peroxisomal abundance similar to that of Pex37p. It is tempting to speculate that based on the similar phenotype and the affiliation to the same protein family that Sym1p is the Pex37p homologue in *S. cerevisiae*. However, additional studies are required, to support this assumption.

We chose to focus on the *pex34*Δ strain that displayed a reduced number of peroxisomes upon *de novo* synthesis conditions (**Supplementary table 1, FIG 3**). Originally, Pex34p was identified by a large-scale protein interaction screen that identified the protein as a binding partner of a number of peroxisomal biogenesis factors (Yu et al (2008)). Later on, it was discovered that Pex34p functions together with Pex11p family members in peroxisome biogenesis (Tower et al., 2011), originally based on the 2-hybrid interactions between Pex34p and the Pex11p family members (Pex11p, Pex25p and Pex27p). More recently, Pex34p was characterized as a tether protein involved in peroxisome-mitochondria (PerMit) contacts and it was shown to contribute to β-oxidation of fatty acids (Shai et al., 2018). Our data suggest that Pex34p also performs a role in the *de novo* synthesis of peroxisomes, which may or may not be due to its role in mediating contact with mitochondria. In this respect, it is interesting to note that members of the Pex11p-family that interact with Pex34p are also involved in the *de novo* formation of peroxisomes (Opalinski et al., 2011). Our results support the notion of a requirement of Pex34p for an efficient exit of Pex3p from the ER. Accordingly, reintroduction of Pex3p-GFP in *pex34*Δ*pex3*Δ cells did not completely restore the formation of peroxisomes (**FIG 5**). When expressed in *pex3*Δ cells, the Pex3p-fusion protein completely localized to peroxisomes, which form *de novo* and proliferate normally under inducing conditions so that the complemented cells are characterized by at least a few peroxisomes. In contrast, when expressed in *pex3*Δ*pex34*Δ cells, the Pex3p-GFP at least partly retains in structures, which might be the ER, and its expression leads to the formation of mostly one peroxisome per cell. This finding might be explained by different possibilities namely 1) defect in the fission of peroxisomes 2) defect in the *de novo* formation of peroxisomes. However, a sole defect in fission of peroxisomes might be difficult to reconcile with the possible ER-retention of the membrane marker Pex3p, unless the defect might feedback and slow down the *de novo* synthesis. With respect to a possible function for Pex34p in the formation of peroxisomes from the ER, it is tempting to speculate that the observed phenotype of mostly one peroxisome per cell might be explained by an inefficient budding of newly formed peroxisomes from the ER. The budding defect is accompanied by the observed retention of Pex3p at the ER in the absence Pex34p, which might be the cause or the consequence of the defect. The fact that Pex3p under these circumstances is distributed throughout the ER might be explained by a role for Pex34p in Pex3p recruitment to the budding zone.

As Pex25p has also been implicated in *de novo* synthesis of peroxisomes in *H. polymorpha* (Saraya et al., 2011) and *S. cerevisiae* (Huber et al., 2012), and as Pex25p and Pex34p do interact, we investigated whether both proteins function in concert in the *de novo* formation of peroxisomes and studied this process in the *pex34*Δ*pex25*Δ double mutant. Cells concurrently lacking Pex25p and Pex34p are characterized by a severe import defect for the otherwise peroxisomal Mdh2-GFP (Tower et al., 2011) and GFP-SKL under inducing conditions (**FIG 6**). Remarkably, the peroxisome biogenesis defect of the double mutant was not effectively rescued by complementation with either Pex34p or Pex25p reinforcing the non-redundant function of these proteins (**FIG 6**).

The data presented show that Pex34p and Pex25p function in concert in the maintenance of peroxisome abundance and in the de novo synthesis of peroxisomes from the ER. The detailed role of Pex34p functions in these two processes is still unknown. The distribution of Pex3p in the ER upon Pex34p-depletion would be in agreement with the role of a Pex3p-specific ER-recruitment factor. The fact that Pex19p can bridge the interaction of Pex34p with other PMPs is unusual and might be explained by a non-typical Pex19p-Pex34p interaction, which needs to be further investigated.

## Acknowledgements

This work was supported by grants of the Deutsche Forschungsgemeinschaft to R. E. (FOR1905), an ERC CoG to M.S (Peroxisystem 646604). M.S is an incumbent of the Dr. Gilbert Omenn and Martha Darling Professorial Chair in Molecular Genetics.

## Notes

### Competing Interest Statement

The authors have declared no competing interest.

## References

Antonenkov VD, Isomursu A, Mennerich D, Vapola MH, Weiher H, Kietzmann T, Hiltunen JK (2015) The Human Mitochondrial DNA Depletion Syndrome Gene MPV17 Encodes a Non-selective Channel That Modulates Membrane Potential. J Biol Chem 290: 13840–13861

Aranovich A, Hua R, Rutenberg AD, Kim PK (2014) PEX16 contributes to peroxisome maintenance by constantly trafficking PEX3 via the ER. Journal of cell science 127: 3675–86

Ausubel FJ, Brent R, Kingston RE, Moore DD, Seidman JG, Smith JA, Struhl K (1992) Current Protocols in Molecular Biology. Greene Publishing Associates, New York

Baldwin HA, Wang C, Kanfer G, Shah HV, Velayos-Baeza A, Dulovic-Mahlow M, Brüggemann N, Anding A, Baehrecke EH, Maric D, Prinz WA, Youle RJ (2021) VPS13D promotes peroxisome biogenesis. J Cell Biol 220: e202001188

Braun A, Kammerer S, Weissenhorn W, Weiss EH, Cleve H (1994) Sequence of a putative human housekeeping gene *(HK33)* localized on chromosome 1. Gene 146: 291–295

Breslow DK, Cameron DM, Collins SR, Schuldiner M, Stewart-Ornstein J, Newman HW, Braun S, Madhani HD, Krogan NJ, Weissman JS (2008) A comprehensive strategy enabling high-resolution functional analysis of the yeast genome. Nat Methods 5: 711–718

Brocard C, Lametschwandtner G, Koudelka R, Hartig A (1997) Pex14p is a member of the protein linkage map of Pex5p. EMBO J 16: 5491–5500

Cepińska MN, Veenhuis M, van der Klei IJ, Nagotu S (2011) Peroxisome fission is associated with reorganization of specific membrane proteins. Traffic 12: 925–937

Chalermwat C, Thosapornvichai T, Wongkittichote P, Phillips JD, Cox JE, Jensen AN, Wattanasirichaigoon D, Jensen LT (2019) Overexpression of the peroxin Pex34p suppresses impaired acetate utilization in yeast lacking the mitochondrial aspartate/glutamate carrier Agc1p. FEMS Yeast Res 19: foz078

Chen Y, Pieuchot L, Loh RA, Yang J, Kari TM, Wong JY, Jedd G (2014) Hydrophobic handoff for direct delivery of peroxisome tail-anchored proteins. Nat Commun 5: doi: 10.1038/ncomms6790.

Chevray PM, Nathans D (1992) Protein interaction cloning in yeast: identification of mammalian proteins that react with the leucine zipper of Jun. Proc Natl Acad Sci USA 89: 5789–5793

Cohen Y, Schuldiner M (2011) Advanced methods for high-throughput microscopy screening of genetically modified yeast libraries. Methods Mol Biol 781: 127–159

Debelyy MO, Platta HW, Saffian D, Hensel A, Thoms S, Meyer HE, Warscheid B, Girzalsky W, Erdmann R (2011) Ubp15p, an ubiquitin hydrolase associated with the peroxisomal export machinery. J Biol Chem 286: 28223–28234

Distel B, Erdmann R, Gould SJ, Blobel G, Crane DI, Cregg JM, Dodt G, Fujiki Y, Goodman JM, Just WW, Kiel JAKW, Kunau W-H, Lazarow PB, Mannaerts GP, Moser HW, Osumi T, Rachubinski RA, Roscher A, Subramani S, Tabak HF et al. (1996) A unified nomenclature for peroxisome biogenesis factors. J Cell Biol 135: 1–3

Eberhart T, Kovacs WJ (2018) Pexophagy in yeast and mammals: an update on mysteries. Histochem Cell Biol 150: 473–488

Erdmann R, Blobel G (1995) Giant peroxisomes in oleic acid-induced *Saccharomyces cerevisiae* lacking the peroxisomal membrane protein Pmp27p. J Cell Biol 128: 509–523

Erdmann R, Kunau W-H (1992) A Genetic Approach to the Biogenesis of Peroxisomes in the Yeast *Saccharomyces cerevisiae*. Cell Biochem Funct 10: 167–174

Erdmann R, Veenhuis M, Mertens D, Kunau W-H (1989) Isolation of peroxisome-deficient mutants of *Saccharomyces cerevisiae*. Proc Natl Acad Sci USA 86: 5419–5423

Fang Y, Morrell JC, Jones JM, Gould SJ (2004) PEX3 functions as a PEX19 docking factor in the import of class I peroxisomal membrane proteins. J Cell Biol 164: 863–875

Fields S, Sternglanz R (1994) The two-hybrid system: an assay for protein-protein interactions. Trends Genet 10:286–292

Gabay-Maskit S, Cruz-Zaragoza LD, Shai N, Eisenstein M, Bibi C, Cohen N, Hansen T, Yifrach E, Harpaz H, Belostotsky R, Schliebs W, Schuldiner M, Erdmann R, Zalckvar E (2020) A piggybacking mechanism enables peroxisomal localization of the glyoxylate cycle enzyme Mdh2 in yeast. J Cell Sci 133: cs244376.

Giaever G, Chu AM, Ni L, Connelly C, Riles L, Véronneau S, et a (2002) Functional profiling of the Saccharomyces cerevisiae genome. Nature 418: 387–391

Gietz RD, Woods RA (1994) High Efficiency transformation in Yeast. In; Molecular Genetics of Yeast: Practical Approaches, ed JA Johnston, Oxford University Press: 121–134

Girzalsky W, Hoffmann LS, Schemenewitz A, Nolte A, Kunau WH, Erdmann R (2006) Pex19p-dependent targeting of Pex17p, a peripheral component of the peroxisomal protein import machinery. J Biol Chem 281: 19417–19425

Girzalsky W, Rehling P, Stein K, Kipper J, Blank L, Kunau W-H, Erdmann R (1999) Involvement of Pex13p in Pex14p localization and peroxisomal targeting signal 2 dependent protein import into peroxisomes. J Cell Biol 144: 1151–1162

Götte K, Girzalsky W, Linkert M, Baumgart E, Kammerer S, Kunau W-H, Erdmann R (1998) Pex19p, a farnesylated protein essential for peroxisome biogenesis. Mol Cell Biol 18: 616–628

Guillén-Samander A, Leonzino M, Hanna MG, Tang N, Shen H, De Camilli P (2021) VPS13D bridges the ER to mitochondria and peroxisomes via Miro. J Cell Biol 220: e202010004

Güldener U, Heck S, Fiedler T, Beinhauer J, Hegemann JH (1996) A new efficient gene disruption cassette for repeated use in budding yeast. Nucleic Acids Res 24: 2519–2524

Haan GJ, Baerends RJ, Krikken AM, Otzen M, Veenhuis M, Klei IJ (2006) Reassembly of peroxisomes in *Hansenula polymorpha* pex3 cells on reintroduction of Pex3p involves the nuclear envelope. FEMS Yeast Res 6: 186–194

Halbach A, Rucktaschel R, Rottensteiner H, Erdmann R (2009) The N-domain of Pex22p can functionally replace the Pex3p N-domain in targeting and peroxisome formation. J Biol Chem 284: 3906–16

Heiland I, Erdmann R (2005) Topogenesis of peroxisomal proteins does not require a functional cytoplasm-to-vacuole transport. Eur J Cell Biol 84: 799–807

Hettema EH, Girzalsky W, van Den Berg M, Erdmann R, Distel B (2000) *Saccharomyces cerevisiae* Pex3p and Pex19p are required for proper localization and stability of peroxisomal membrane proteins. EMBO J 19: 223–233

Hettema EH, Motley AM (2009) How peroxisomes multiply. J Cell Sci 15: 2331–2336

Hoepfner D, Schildknegt D, Braakman I, Philippsen P, Tabak HF (2005) Contribution of the endoplasmic reticulum to peroxisome formation. Cell 122: 89–95

Hoepfner D, van den Berg M, Philippsen P, Tabak HF, Hettema EH (2001) A role for Vps1p, actin, and the Myo2p motor in peroxisome abundance and inheritance in *Saccharomyces cerevisiae*. J Cell Biol 155: 979–990

Huber A, Koch J, Kragler F, Brocard C, Hartig A (2012) A subtle interplay between three Pex11 proteins shapes de novo formation and fission of peroxisomes. Traffic 13: 157–167

Islinger M, Cardoso MJ, Schrader M (2010) Be different - the diversity of peroxisomes in the animal kingdom. Biochim Biophys Acta 1803: 881–897

Islinger M, Voelkl A, Fahimi HD, Schrader M (2018) The peroxisome: an update on mysteries 2.0. Histochem Cell Biol 150: 443–471

James GL, Goldstein JL, Pathak RK, Anderson RGW, Brown MS (1994) PxF, a prenylated protein of peroxisomes. J Biol Chem 269: 14182–14190

Jansen RLM, van der Klei IJ (2019) The peroxisome biogenesis factors Pex3 and Pex19: multitasking proteins with disputed functions. FEBS Lett 593: 457–474

Jones JM, Morrell JC, Gould SJ (2004) PEX19 is a predominantly cytosolic chaperone and import receptor for class 1 peroxisomal membrane proteins. J Cell Biol 164: 57–67

Kammerer S, Arnold N, Gutensohn W, Mewes HW, Kunau WH, Hofler G, Roscher AA, Braun A (1997) Genomic organization and molecular characterization of a gene encoding HsPXF, a human peroxisomal farnesylated protein. Genomics 45: 200–210

Kataya ARA, Muench DG, Moorhead GB (2019) A Framework to Investigate Peroxisomal Protein Phosphorylation in Arabidopsis. Trends Plant Sci 24: 366–381

Kerssen D, Hambruch E, Klaas W, Platta HW, de Kruijff B, Erdmann R, Kunau WH, Schliebs W (2006) Membrane association of the cycling peroxisome import receptor Pex5p. J Biol Chem 281: 27003–27015

Kragt A, Voorn-Brouwer T, van den Berg M, Distel B (2005) Endoplasmic reticulum-directed Pex3p routes to peroxisomes and restores peroxisome formation in a Saccharomyces cerevisiae pex3Delta strain. J Biol Chem 280: 34350–7

Kunze M, Pracharoenwattana I, Smith SM, Hartig A (2006) A central role for the peroxisomal membrane in glyoxylate cycle function. Biochim Biophys Acta 1763: 1441–1452

Kuravi K, Nagotu S, Krikken AM, Sjollema K, Deckers M, Erdmann R, Veenhuis M, van der Klei IJ (2006) Dynamin-related proteins Vps1p and Dnm1p control peroxisome abundance in Saccharomyces cerevisiae. Journal of cell science 119: 3994–4001

Lillo C, Kataya ARA, Heidari B, Creighton HT, Nemie-Feyissa D, Ginbot Z, Jonassen EM (2014) Protein phosphatases PP2A, PP4 and PP6: mediators and regulators in development and responses to environmental cues. Plant Cell Environ 37: 2631–3648

Lygerou Z, Christophides G, Séraphin B (1999) A Novel Genetic Screen for snRNP Assembly Factors in Yeast Identifies a Conserved Protein, Sad1p, Also Required for Pre-mRNA Splicing. Mol Cell Biol 19: 2008–2020

Ma C, Agrawal G, Subramani S (2011) Peroxisome assembly: matrix and membrane protein biogenesis. J Cell Biol 193: 7–16

Matsuzono Y, Fujiki Y (2006) In vitro transport of membrane proteins to peroxisomes by shuttling receptor Pex19p. J Biol Chem 281: 36–42

Motley AM, Hettema EH (2007) Yeast peroxisomes multiply by growth and division. J Cell Biol 178: 399–410

Motley AM, Ward GP, Hettema EH (2008) Dnm1p-dependent peroxisome fission requires Caf4p, Mdv1p and Fis1p. J Cell Sci 121: 1633–40

Niedenthal RK, Riles L, Johnston M, Hegemann JH (1996) Green fluorescent protein as a marker for gene expression and subcellular localization in budding yeast. Yeast 12: 773–786

Nordgren M, Wang B, Apanasets O, Fransen M Peroxisome degradation in mammals: mechanisms of action, recent advances, and perspectives. Front Physiol 4: doi: 10.3389/fphys.2013.00145

Opalinski L, Kiel JA, Williams C, Veenhuis M, van der Klei IJ (2011) Membrane curvature during peroxisome fission requires Pex11. EMBO J 30: 5–16

Papa FR, Hochstrasser M (1993) The yeast DOA4 gene encodes a deubiquitinating enzyme related to a product of the human tre-2 oncogene. Nature 366: 313–9

Purdue PE, Lazarow PB (2001) Pex18p is constitutively degraded during peroxisome biogenesis. J Biol Chem 276: 47684–47689

Rehling P, Marzioch M, Niesen F, Wittke E, Veenhuis M, Kunau W-H (1996b) The import receptor for the peroxisomal targeting signal 2 (PTS2) in *Saccharomyces cerevisiae* is encoded by the *PAS7* gene. EMBO J 15: 2901–2913

Rokka A, Antonenkov VD, Soininen R, Immonen HL, Pirilä PL, Bergmann U, Sormunen RT, Weckström M, Benz R, Hiltunen JK (2009) Pxmp2 is a channel-forming protein in Mammalian peroxisomal membrane. PloS one 4: e5090

Rottensteiner H, Kramer A, Lorenzen S, Stein K, Landgraf C, Volkmer-Engert R, Erdmann R (2004) Peroxisomal membrane proteins contain common Pex19p-binding sites that are an integral part of their targeting signals (mPTS). Mol Biol Cell 7: 3406–3417

Rottensteiner H, Stein K, Sonnenhol E, Erdmann R (2003) Conserved function of pex11p and the novel pex25p and pex27p in peroxisome biogenesis. Mol Biol Cell 14: 4316–4328

Saleem RA, Knoblach B, Mast FD, Smith JJ, Boyle J, Dobson CM, Long-O’Donnell R, Rachubinski RA, Aitchison JD (2008) Genome-wide analysis of signaling networks regulating fatty acid-induced gene expression and organelle biogenesis. J Cell Biol 181: 281–92

Saraya R, Krikken AM, Veenhuis M, van der Klei IJ (2011) Peroxisome reintroduction in Hansenula polymorpha requires Pex25 and Rho1. J Cell Biol 193: 885–900

Schrader M, Pellegrini L (2017) The making of a mammalian peroxisome, version 2.0: mitochondria get into the mix. Cell Death Differ 24: 1148–1152

Schummer A, Maier R, Gabay-Maskit S, Hansen T, Mühlhäuser WWD, Suppanz I, Fadel A, Schuldiner M, Girzalsky W, Oeljeklaus S, Zalckvar E, Erdmann R, Warscheid B (2020) Pex14p Phosphorylation Modulates Import of Citrate Synthase 2 Into Peroxisomes in Saccharomyces cerevisiae. Front Cell Dev Biol 15: 549451

Shai N, Yifrach E, van Roermund CWT, Cohen N, Bibi C, IJlst L, Cavellini L, Meurisse J, Schuster R, Zada L, Mari MC, Reggiori FM, Hughes AL, Escobar-Henriques M, Cohen MM, Waterham HR, Wanders RJA, Schuldiner M, Zalckvar E (2018) Systematic mapping of contact sites reveals tethers and a function for the peroxisome-mitochondria contact. Nat Commun 9: 1761

Shibata H, Kashiwayama Y, Imanaka T, Kato H (2004) Domain architecture and activity of human Pex19p, a chaperone-like protein for intracellular trafficking of peroxisomal membrane proteins. J Biol Chem 279: 38486–38494

Singh R, Manivannan S, Krikken AM, de Boer R, Bordin N, Devos DP, van der Klei IJ (2019) Hansenula polymorpha Pex37 is a peroxisomal membrane protein required for organelle fission and segregation. FEBS J 287: 1742–1757

Smith JJ, Marelli M, Christmas RH, Vizeacoumar FJ, Dilworth DJ, Ideker T, Galitski T, Dimitrov K, Rachubinski RA, Aitchison JD (2002) Transcriptome profiling to identify genes involved in peroxisome assembly and function. J Cell Biol 158: 259–271

Tam YY, Rachubinski RA (2002) *Yarrowia lipolytica* Cells Mutant for the *PEX24* Gene Encoding a Peroxisomal Membrane Peroxin Mislocalize Peroxisomal Proteins and Accumulate Membrane Structures Containing Both Peroxisomal Matrix and Membrane Proteins. Mol Biol Cell 13: 2681–2691

Tam YY, Torres-Guzman JC, Vizeacoumar FJ, Smith JJ, Marelli M, Aitchison JD, Rachubinski RA (2003) Pex11-related Proteins in Peroxisome Dynamics: A Role for the Novel Peroxin Pex27p in Controlling Peroxisome Size and Number in *Saccharomyces cerevisiae*. Mol Biol Cell 14: 4098–4102

Titorenko VI, Ogrydziak DM, Rachubinski RA (1997) Four distinct secretory pathways serve protein secretion, cell surface growth, and peroxisome biogenesis in the yeast *Yarrowia lipolytica*. Mol Cell Biol 17: 5210–5226

Tong AH, Boone C (2006) Synthetic genetic array analysis in Saccharomyces cerevisiae. Methods Mol Biol 313: 171–192

Tower RJ, Fagarasanu A, Aitchison JD, Rachubinski RA (2011) The peroxin Pex34p functions with the Pex11 family of peroxisomal divisional proteins to regulate the peroxisome population in yeast. Mol Biol Cell 22: 1727–1738

Trott A, Morano KA (2004) SYM1 is the stress-inducedSaccharomyces cerevisiae ortholog of the mammalian kidney diseasegene Mpv17 and is required for ethanol metabolism and tolerance duringheat shock. Eukaryot Cell 3: 620–631

van der Zand A, Braakman I, Tabak HF (2010) Peroxisomal membrane proteins insert into the endoplasmic reticulum. Mol Biol Cell 21: 2057–2065

Vizeacoumar FJ, Torres-Guzman JC, Tam YY, Aitchison JD, Rachubinski RA (2003) YHR150w and YDR479c encode peroxisomal integral membrane proteins involved in the regulation of peroxisome number, size, and distribution in *Saccharomyces cerevisiae*. J Cell Biol 161: 321–332

Walter T, Erdmann R (2019) Current Advances in Protein Import into Peroxisomes. Protein J 38: 351–362

Wróblewska JP, van der Klei IJ (2019) Peroxisome Maintenance Depends on De Novo Peroxisome Formation in Yeast Mutants Defective in Peroxisome Fission and Inheritance. Int J Mol Sci 20: pii: E4023. doi: 10.3390/ijms20164023.

Yuan W, Veenhuis M, van der Klei IJ (2016) The birth of yeast peroxisomes. Biochim Biophys Acta 1863: 902–910

