## Supplemental Table 1 for "Identification of Pex34p as a component of the peroxisomal de novo biogenesis machinery in yeast"

### Supplementary Table 1 – List of hits from the screen

C – cytosolic distribution of the reporter; S – small peroxisomes; L – large peroxisomes; R – Reduced number of peroxisomes; El – elongated peroxisomes; M – mixes phenotype

| ORF | Gene | Phenotype | Overlapping genes in case of dubious ORFS | SGD Description |
|---|---|---|---|---|
| *YAL055W* | *PEX22* | C | | Putative peroxisomal membrane protein; required for import of peroxisomal proteins; functionally complements a Pichia pastoris pex22 mutation |
| *YDR244W* | *PEX5* | C | | Peroxisomal membrane signal receptor for peroxisomal matrix proteins; receptor for the C-terminal tripeptide signal sequence (PTS1) of peroxisomal matrix proteins; required for peroxisomal matrix protein import; also proposed to have PTS1-receptor independent functions |
| *YDR329C* | *PEX3* | C | | Peroxisomal membrane protein (PMP); required for proper localization and stability of PMPs; anchors peroxisome retention factor Inp1p at the peroxisomal membrane; interacts with Pex19p |
| *YGL152C* | *YGL152C* | C | Pex14: Central component of the peroxisomal importomer complex; peroxisomal protein import machinery docking complex component; interacts with both PTS1 (Pex5p) and PTS2 (Pex7p) peroxisomal matrix protein signal recognition factors and membrane receptor Pex13p | Dubious open reading frame; unlikely to encode a functional protein, based on available experimental and comparative sequence data; partially overlaps the verified ORF PEX14/YGL153W |
| *YGL153W* | *PEX14* | C | | Central component of the peroxisomal importomer complex; peroxisomal protein import machinery docking complex component; interacts with both PTS1 (Pex5p) and PTS2 (Pex7p) peroxisomal matrix protein signal recognition factors and membrane receptor Pex13p |
| *YGR077C* | *PEX8* | C | | Intraperoxisomal organizer of the peroxisomal import machinery; organizes the formation of the importomer complex, bridging the docking complex with the RING finger complex; tightly associated with the lumenal face of the peroxisomal membrane; essential for peroxisome biogenesis; binds PTS1-signal receptor Pex5p, and PTS2-signal receptor Pex7p |
| *YGR133W* | *PEX4* | C | | Peroxisomal ubiquitin conjugating enzyme; required for peroxisomal matrix protein import and peroxisome biogenesis |
| *YJL210W* | *PEX2* | C | | RING-finger peroxin and E3 ubiquitin ligase; peroxisomal membrane protein with a C-terminal zinc-binding RING domain, forms translocation subcomplex with Pex10p and Pex12p which functions in peroxisomal matrix protein import |
| *YJL211C* | *YJL211C* | C | Pex2: RING-finger peroxin and E3 ubiquitin ligase; peroxisomal membrane protein with a C-terminal zinc-binding RING domain, forms translocation subcomplex with Pex10p and Pex12p which functions in peroxisomal matrix protein import | Dubious open reading frame; unlikely to encode a functional protein, based on available experimental and comparative sequence data; partially overlaps the verified gene YJL210W/PEX2 |

| | | | | |
|---|---|---|---|---|
| *YLR191W* | *PEX13* | C | | Peroxisomal importomer complex component; integral peroxisomal membrane protein required for docking and translocation of peroxisomal matrix proteins; interacts with the PTS1 signal recognition factor Pex5p and the PTS2 signal recognition factor Pex7p; forms a complex with Pex14p and Pex17p; human homolog PEX13 complements yeast null mutant |
| *YMR026C* | *PEX12* | C | | C3HC4-type RING-finger peroxin and E3 ubiquitin ligase; required for peroxisome biogenesis and peroxisomal matrix protein import; forms translocation subcomplex with Pex2p and Pex10p; mutations in human homolog cause peroxisomal disorder |
| *YNL214W* | *PEX17* | C | | Membrane peroxin of the peroxisomal importomer complex; complex facilitates the import of peroxisomal matrix proteins; required for peroxisome biogenesis |
| *YNL329C* | *PEX6* | C | | AAA-peroxin; heterodimerizes with AAA-peroxin Pex1p and participates in the recycling of peroxisomal signal receptor Pex5p from the peroxisomal membrane to the cystosol; mutations in human PEX6 can lead to severe peroxisomal disorders and early death |
| *YOL044W* | *PEX15* | C | | Tail-anchored type II integral peroxisomal membrane protein; required for peroxisome biogenesis; cells lacking Pex15p mislocalize peroxisomal matrix proteins to cytosol; overexpression results in impaired peroxisome assembly |
| *YPL172C* | *COX10* | C | | Heme A:farnesyltransferase; catalyzes first step in conversion of protoheme to heme A prosthetic group required for cytochrome c oxidase activity; human ortholog COX10 can complement yeast cox10 null mutant; human ortholog COX10 is associated with mitochondrial disorders |
| *YBR187W* | *GDT1* | C, R | | Calcium and manganese transporter with higher affinity for Ca2+; involved in Ca2+ and Mn2+ homeostasis; localizes to the cis- and medial-Golgi apparatus; GFP-fusion localizes to the vacuole; required for the Mn2+-dependent function of glycosylation enzymes; TMEM165, a human ortholog linked to Congenital Disorders of Glycosylation, functionally complements the null allele; expression pattern and physical interactions suggest a possible role in ribosome biogenesis; expression regulated by Gcr1p |
| *YCL056C* | *PEX34* | C, R | | Protein that regulates peroxisome populations; peroxisomal integral membrane protein; interacts with Pex11p, Pex25p, and Pex27p to control both constitutive peroxisome division and peroxisome morphology and abundance during peroxisome proliferation |
| *YDR080W* | *VPS41* | C, R | | Subunit of the HOPS endocytic tethering complex; vacuole membrane protein that functions as a Rab GTPase effector, interacting specifically with the GTP-bound conformation of Ypt7p, facilitating tethering, docking and promoting membrane fusion events at the late endosome and vacuole; required for both membrane and protein trafficking; Yck3p-mediated phosphorylation regulates the organization of vacuolar fusion sites |
| *YDR265W* | *PEX10* | C, R | | Peroxisomal membrane E3 ubiquitin ligase; required for for Ubc4p-dependent Pex5p ubiquitination and peroxisomal matrix protein import; contains zinc-binding RING domain; mutations in human homolog cause various peroxisomal disorders |
| *YDR424C* | *GAL7* | C, R | | Cytoplasmic light chain dynein, microtubule motor protein; required for intracellular transport and cell division; involved in mitotic spindle positioning; forms complex with dynein intermediate chain Pac11p that promotes Dyn1p homodimerization, potentiates motor processivity; Dyn2p-Pac11p complex important for interaction of dynein motor complex with dynactin complex; acts as molecular glue to dimerize, stabilize Nup82-Nsp1-Nup159 complex module of cytoplasmic pore filaments |
| *YGL212W* | *VAM7* | C, R | | Vacuolar SNARE protein; functions with Vam3p in vacuolar protein trafficking; has an N-terminal PX domain (phosphoinositide-binding module) that binds PtdIns-3-P and mediates membrane binding; SNAP-25 homolog; protein abundance increases in response to DNA replication stress |
| *YIL035C* | *CKA1* | C, R | | Alpha catalytic subunit of casein kinase 2 (CK2); a Ser/Thr protein kinase with roles in cell growth and proliferation; CK2, comprised of CKA1, CKA2, CKB1 and CKB2, has many substrates including transcription factors and all RNA polymerases; regulates Fkh1p-mediated donor preference during mating-type switching |
| *YKL197C* | *PEX1* | C, R | | AAA-peroxin; heterodimerizes with AAA-peroxin Pex6p and participates in the recycling of peroxisomal signal receptor Pex5p from the peroxisomal membrane to the cystosol; induced by oleic acid and upregulated during anaerobiosis; mutations in human PEX1 can lead to severe peroxisomal disorders and early death |
| *YMR288W* | *HSH155* | C, R | | U2-snRNP associated splicing factor; forms extensive associations with the branch site-3' splice site-3' exon region upon prespliceosome formation; similarity to the mammalian U2 snRNP-associated splicing factor SAP155 |
| *YNL121C* | *TOM70* | C, R | | Component of the TOM (translocase of outer membrane) complex; involved in the recognition and initial import steps for all mitochondrially directed proteins; acts as a receptor for incoming precursor proteins; TOM70 has a paralog, TOM71, that arose from the whole genome duplication |
| *YOR060C* | *SLD7* | C, R | | Protein with a role in chromosomal DNA replication; interacts with Sld3p and reduces its affinity for Cdc45p; deletion mutant has aberrant mitochondria; ortholog of human MTBP, which is a DNA replication origin firing factor |
| *YPL112C* | *PEX25* | C, R | | Peripheral peroxisomal membrane peroxin; required for the regulation of peroxisome size and maintenance, recruits GTPase Rho1p to peroxisomes, induced by oleate, interacts with Pex27p; PEX25 has a paralog, PEX27, that arose from the whole genome duplication |

| | | | | |
|---|---|---|---|---|
| *YDL099W* | *BUG1* | EL | | Cis-golgi localized protein involved in ER to Golgi transport; forms a complex with the mammalian GRASP65 homolog, Grh1p; mutants are compromised for the fusion of ER-derived vesicles with Golgi membranes |
| *YAL016C-B* | *YAL016C-B* | M | Most likely interrupts Psk1: PAS domain-containing serine/threonine protein kinase; coordinately regulates protein synthesis and carbohydrate metabolism and storage in response to a unknown metabolite that reflects nutritional status | Dubious open reading frame; unlikely to encode a functional protein, based on available experimental and comparative sequence data |
| *YAL027W* | *SAW1* | M | | 5'- and 3'-flap DNA binding protein; recruits Rad1p-Rad10p to single-strand annealing intermediates with 3' non-homologous tails for removal during double-strand break repair; complexes with Rad1p-Rad10p and stimulates its endonuclease activity; green fluorescent protein (GFP)-fusion protein localizes to the nucleus |
| *YAL037C-A* | *YAL037C-A* | M | | Putative protein of unknown function |
| *YAR035C-A* | *YAR035C-A* | M | | Putative protein of unknown function; emerging ORF that arose de novo from non-genic locus; identified by gene-trapping, microarray-based expression analysis, and genome-wide homology searching; localizes to mitochondria |
| *YBR178W* | *YBR178W* | M | Overlaps Eht1: Octanoyl-CoA:ethanol acyltransferase; also functions as thioesterase; plays a minor role in medium-chain fatty acid ethyl ester biosynthesis | Dubious open reading frame; unlikely to encode a functional protein, based on available experimental and comparative sequence data; partially overlaps the verified gene YBR177C |
| *YBR183W* | *YBR183W* | M | | Alkaline ceramidase; also has reverse (CoA-independent) ceramide synthase activity; catalyzes both breakdown and synthesis of phytoceramide; overexpression confers fumonisin B1 resistance; YPC1 has a paralog, YDC1, that arose from the whole genome duplication |
| *YBR184W* | *YBR184W* | M | | Putative protein of unknown function; YBR184W is not an essential gene |
| *YBR186W* | *PCH2* | M | | Hexameric ring ATPase that remodels chromosome axis protein Hop1p; nucleolar component of the pachytene checkpoint, which prevents chromosome segregation when recombination and chromosome synapsis are defective; also represses meiotic interhomolog recombination in rDNA; required for meiotic double-stranded break formation |
| *YBR200W-A* | *YBR200W-A* | M | | Putative protein of unknown function; identified by fungal homology and RT-PCR |
| *YDR476C* | *YDR476C* | M | | Putative protein of unknown function; green fluorescent protein (GFP)-fusion protein localizes to the endoplasmic reticulum; YDR476C is not an essential gene |
| *YFL001W* | *DEG1* | M | | tRNA:pseudouridine synthase; introduces pseudouridines at position 38 or 39 in tRNA; also responsible for pseudouracil modification of some mRNAs; important for maintenance of translation efficiency and normal cell growth, localizes to both the nucleus and cytoplasm; non-essential for viability |
| *YGL116W* | *CDC20* | M | | Activator of anaphase-promoting complex/cyclosome (APC/C); APC/C is required for metaphase/anaphase transition; directs ubiquitination of mitotic cyclins, Pds1p, and other anaphase inhibitors; cell-cycle regulated; potential Cdc28p substrate; relative distribution to the nucleus increases upon DNA replication stress |
| *YGL124C* | *MON1* | M | | Subunit of a heterodimeric guanine nucleotide exchange factor (GEF); subunit of the Mon1-Ccz1 GEF complex which stimulates nucleotide exchange and activation of Ypt7p, a Rab family GTPase involved in membrane tethering and fusion events at the late endosome and vacuole; GEF activity is stimulated by membrane association and anionic phospholipids; role in localizing Ypt7p to the vacuolar membrane; required for autophagy, the CVT pathway and mitophagy; potential Cdc28 substrate |

| | | | | |
|---|---|---|---|---|
| *YGL169W* | *SUA5* | M | | Protein involved in threonylcarbamoyl adenosine biosynthesis; Sua5p and Qri7p are necessary and sufficient for RNA t6A modification in vitro; null mutant lacks N6-threonylcarbamoyl adenosine (t6A) modification in the anticodon loop of ANN-decoding tRNA; member of conserved YrdC/Sua5 family; binds single-stranded telomeric DNA and null mutant has abnormal telomere length |
| *YGR269W* | *YGR269W* | M | Hua1: Cytoplasmic protein containing a zinc finger domain; sequence similarity to that of Type I J-proteins | Dubious open reading frame; unlikely to encode a functional protein, based on available experimental and comparative sequence data; partially overlaps the uncharacterized ORF HUA1/YGR268C |
| *YIL148W* | *RPL40A* | M | | Ubiquitin-ribosomal 60S subunit protein L40A fusion protein; cleaved to yield ubiquitin and ribosomal protein L40A; ubiquitin may facilitate assembly of the ribosomal protein into ribosomes; homologous to mammalian ribosomal protein L40, no bacterial homolog; RPL40A has a paralog, RPL40B, that arose from the whole genome duplication; relative distribution to the nucleus increases upon DNA replication stress |
| *YIR033W* | *MGA2* | M | | ER membrane protein involved in regulation of OLE1 transcription; inactive ER form dimerizes and one subunit is then activated by ubiquitin/proteasome-dependent processing followed by nuclear targeting; MGA2 has a paralog, SPT23, that arose from the whole genome duplication |
| *YKL213C* | *DOA1* | M | | WD-repeat protein involved in ubiquitin-mediated protein degradation; Ub-binding protein with a role in Ub homeostasis; substrate-recruiting adaptor for Cdc48p in mitochondria-associated degradation; inhibits degradation of Ufd2p-dependent substrates; facilitates proteolysis of Cse4p, a centromeric H3-like protein; required for ribophagy; promotes NHEJ in postdiauxic/stationary phase; protein increases in abundance and relocalizes from nucleus to nuclear periphery upon DNA replication stress |
| *YLL040C* | *VPS13* | M | | Protein involved in prospore membrane morphogenesis; peripheral membrane protein that localizes to the prospore membrane and at numerous membrane contact sites; involved in sporulation, vacuolar protein sorting, prospore membrane formation during sporulation, and protein-Golgi retention; required for mitochondrial integrity; contains a PH-like domain; homologous to human CHAC and COH1 which are involved in Chorea-acanthocytosis and Cohen syndrome, respectively |
| *YLL063C* | *AYT1* | M | | Acetyltransferase; catalyzes trichothecene 3-O-acetylation, suggesting a possible role in trichothecene biosynthesis |
| *YLR166C* | *SEC10* | M | | Essential 100kDa subunit of the exocyst complex; the exocyst mediates polarized targeting and tethering of post-Golgi secretory vesicles to active sites of exocytosis at the plasma membrane prior to SNARE-mediated fusion |
| *YLR305C* | *STT4* | M | | Phosphatidylinositol-4-kinase; functions in the Pkc1p protein kinase pathway; required for normal vacuole morphology, cell wall integrity, and actin cytoskeleton organization; required for autophagosomeâ€“vacuole fusion during autophagy and for lipophagy in both stationary phase cells and during nitrogen starvation; localizes to the plasma membrane and mitochondria in HTP studies |
| *YLR310C* | *CDC25* | M | | Membrane bound guanine nucleotide exchange factor; also known as a GEF or GDP-release factor; indirectly regulates adenylate cyclase through activation of Ras1p and Ras2p by stimulating the exchange of GDP for GTP; required for progression through G1; thermosensitivity of the cdc25-5 mutant is functionally complemented by human RASGRF1 or by a fragment of human SOS1 comprising the CDC25-related catalytic domain |
| *YLR397C* | *AFG2* | M | | ATPase of the CDC48/PAS1/SEC18 (AAA) family, forms a hexameric complex; is essential for pre-60S maturation and release of several preribosome maturation factors; releases Rlp24p from purified pre-60S particles in vitro; target of the ribosomal biosynthesis inhibitor diazaborine; may be involved in degradation of aberrant mRNAs |
| *YML013W* | *UBX2* | M | | Bridging factor involved in ER-associated protein degradation (ERAD); bridges the cytosolic Cdc48p-Npl1p-Ufd1p ATPase complex and the membrane associated Ssm4p and Hrd1p ubiquitin ligase complexes; contains a UBX (ubiquitin regulatory X) domain and a ubiquitin-associated (UBA) domain; redistributes from the ER to lipid droplets during the diauxic shift and stationary phase; required for the maintenance of lipid homeostasis; required for mitochondrial protein translocation-associated degradation |
| *YML079W* | *YML079W* | M | | Non-essential protein of unknown function; has structural resemblance to plant storage and ligand binding proteins (canavalin, glycinin, auxin binding protein) and to some enzymes (epimerase, germin); localizes to the nucleus and cytoplasm |
| *YMR035W* | *IMP2* | M | | Catalytic subunit of mitochondrial inner membrane peptidase complex; required for maturation of mitochondrial proteins of the intermembrane space; complex contains two catalytic subunits (Imp1p and Imp2p that differ in substrate specificity), and Som1p |
| *YMR290C* | *HAS1* | M | | ATP-dependent RNA helicase; involved in the biogenesis of 40S and 60S ribosome subunits; localizes to both the nuclear periphery and nucleolus; highly enriched in nuclear pore complex fractions; constituent of 66S pre-ribosomal particles |

| | | | | |
|---|---|---|---|---|
| *YNL132W* | *KRE33* | M | | Acetyltransferase required for biogenesis of small ribosomal subunit; responsible for incorporation of N4-acetylcytidine into mRNAs; heterozygous mutant shows haploinsufficiency in K1 killer toxin resistance; essential gene; NAT10, the human homolog, implicated in several types of cancer and premature aging |
| *YNL175C* | *NOP13* | M | | Nucleolar protein found in preribosomal complexes; contains an RNA recognition motif (RRM); relative distribution to the nucleolus increases upon DNA replication stress |
| *YNL307C* | *MCK1* | M | | Dual-specificity ser/thr and tyrosine protein kinase; roles in chromosome segregation, meiotic entry, genome stability, phosphorylation-dependent protein degradation (Rcn1p and Cdc6p), inhibition of protein kinase A, transcriptional regulation, inhibition of RNA pol III, calcium stress and inhibition of Clb2p-Cdc28p after nuclear division; MCK1 has a paralog, YGK3, that arose from the whole genome duplication |
| *YOL110W* | *SHR5* | M | | Palmitoyltransferase subunit; this complex adds a palmitoyl lipid moiety to heterolipidated substrates such as Ras1p and Ras2p through a thioester linkage; palmitoylation is required for Ras2p membrane localization; Palmitoyltransferase is composed of Shr5p and Erf2 |
| *YOL126C* | *MDH2* | M | | Cytoplasmic malate dehydrogenase; one of three isozymes that catalyze interconversion of malate and oxaloacetate; involved in the glyoxylate cycle and gluconeogenesis during growth on two-carbon compounds; interacts with Pck1p and Fbp1; mutation in human homolog MDH2 causes early-onset severe encephalopathy |
| *YOR048C* | *RAT1* | M | | Nuclear 5' to 3' single-stranded RNA exonuclease; involved in RNA metabolism, including rRNA and snoRNA processing, as well as poly (A+) dependent and independent mRNA transcription termination; required for cotranscriptional pre-rRNA cleavage; displaces Cdk1p from elongating transcripts, especially as RNAPII reaches the poly(A) site, negatively regulates phosphorylation of the CTD of RNAPII, and inhibits RNAPII transcriptional elongation |
| *YOR350C* | *MNE1* | M | | Protein involved in splicing Group I aI5-beta intron from COX1 mRNA; mitochondrial matrix protein |
| *YPL194W* | *DDC1* | M | | DNA damage checkpoint protein; part of a PCNA-like complex required for DNA damage response, required for pachytene checkpoint to inhibit cell cycle in response to unrepaired recombination intermediates; potential Cdc28p substrate; forms nuclear foci upon DNA replication stress |
| *YPL196W* | *OXR1* | M | | Protein of unknown function required for oxidative damage resistance; required for normal levels of resistance to oxidative damage; null mutants are sensitive to hydrogen peroxide; member of a conserved family of proteins found in eukaryotes |
| *YPL208W* | *RKM1* | M | | SET-domain lysine-N-methyltransferase; catalyzes the formation of dimethyllysine residues on the large ribosomal subunit proteins L23 (Rpl23Ap and Rpl23Bp) and monomethyllysine residues on L18 (Rps18Ap and Rps18Bp) |
| *YER137C* | *YER137C* | M, C | | Protein of unknown function |
| *YDR207C* | *UME6* | R | | Rpd3L histone deacetylase complex subunit; key transcriptional regulator of early meiotic genes; involved in chromatin remodeling and transcriptional repression via DNA looping; binds URS1 upstream regulatory sequence, represses transcription by recruiting conserved histone deacetylase Rpd3p (through co-repressor Sin3p) and chromatin-remodeling factor Isw2p; couples metabolic responses to nutritional cues with initiation and progression of meiosis |
| *YDR376W* | *ARH1* | R | | Oxidoreductase of the mitochondrial inner membrane; involved in cytoplasmic and mitochondrial iron homeostasis and required for activity of Fe-S cluster-containing enzymes; one of the few mitochondrial proteins essential for viability |
| *YKL064W* | *MNR2* | R | | Vacuolar membrane protein required for magnesium homeostasis; putative magnesium transporter; has similarity to Alr1p and Alr2p, which mediate influx of Mg2+ and other divalent cations; localizes to sites of contact between the vacuole and mitochondria (vCLAMPs) |
| *YKR001C* | *VPS1* | R | | Dynamin-like GTPase required for vacuolar sorting; promotes fission of retrograde transport carriers from endosome; also involved in actin cytoskeleton organization, endocytosis, late Golgi-retention of some proteins, regulation of peroxisome biogenesis |
| *YLR151C* | *PCD1* | R | | 8-oxo-dGTP diphosphatase; prevents spontaneous mutagenesis via sanitization of oxidized purine nucleoside triphosphates; can also act as peroxisomal pyrophosphatase with specificity for coenzyme A and CoA derivatives, may function to remove potentially toxic oxidized CoA disulfide from peroxisomes to maintain the capacity for beta-oxidation of fatty acids; nudix hydrolase family member; similar E. coli MutT and human, rat and mouse MTH1 |
| *YLR222C* | *UTP13* | R | | Nucleolar protein; component of the small subunit (SSU) processome containing the U3 snoRNA that is involved in processing of pre-18S rRNA |
| *YLR251W* | *SYM1* | R | | Protein required for ethanol metabolism; induced by heat shock and localized to the inner mitochondrial membrane; homologous to mammalian peroxisomal membrane protein Mpv17; human homolog MPV17 is implicated in hepatocerebral mtDNA depletion syndromes (MDDS), and complements yeast null mutant |
| *YLR422W* | *DCK1* | R | | |

| | | | | |
|---|---|---|---|---|
| *YMR018W* | *PEX9* | R | | Dock family protein (Dedicator Of CytoKinesis), homolog of human DOCK1; upstream component for regulation through the small GTPase Rho5p; may form a complex with Lmo1p that acts as a GEF for Rho5p; interacts with Ino4p; cytoplasmic protein that relocates to mitochondria under oxidative stress; implicated in mitophagy; not an essential protein; DOCK proteins act as guanine nucleotide exchange factors<br>Peroxisomal membrane signal receptor for peroxisomal matrix proteins; oleate-inducible condition-specific import receptor for a subset of PTS1-containing matrix proteins; localizes to both the cytosol and the peroxisomal membrane; similar to human PEX5Rp, a peroxin protein 5 related protein; paralog of Pex5p<br>Alpha-1,6-mannosyltransferase localized to the ER; responsible for addition of alpha-1,6 mannose to dolichol-linked Man7GlcNAc2; acts in the dolichol pathway for N-glycosylation; human homolog ALG12 complements yeast null mutant |
| *YNR030W* | *ALG12* | R | | |
| *YOL138C* | *RTC1* | R | | Subunit of SEACAT, a subcomplex of the SEA complex; Rtc1p, along with Mtc5p and Sea4p, redundantly inhibit the TORC1 inhibitory role of the Iml1p/SEACIT (Iml1p-Npr2p-Npr3p) subcomplex, a GAP for GTPase Gtr1p (EGOC subunit) in response to amino acid limitation, thereby resulting in activation of TORC1 signaling; SEA is a coatomer-related complex that associates dynamically with the vacuole; has N-terminal WD-40 repeats and a C-terminal RING motif; null suppresses cdc13-1<br>Mitochondrial inner membrane localized ATP-dependent DNA helicase; required for the maintenance of the mitochondrial genome; not required for mitochondrial transcription; has homology to E. coli helicase uvrD<br>Mitochondrial protein of unknown function; mCherry fusion protein localizes to the vacuole<br>Galactose-1-phosphate uridyl transferase; synthesizes glucose-1-phosphate and UDP-galactose from UDP-D-glucose and alpha-D-galactose-1-phosphate in the second step of galactose catabolism; human homolog UGP2 can complement yeast null mutant<br>High-mobility group (HMG) protein; binds to and remodels nucleosomes; involved in recruiting FACT and other chromatin remodelling complexes to the chromosomes; functionally redundant with Nhp6Ap; required for transcriptional initiation fidelity of some tRNA genes; homologous to mammalian HMGB1 and HMGB2; NHP6B has a paralog, NHP6A, that arose from the whole genome duplication<br>Subunit of the RNA polymerase II mediator complex; associates with core polymerase subunits to form the RNA polymerase II holoenzyme; essential for transcriptional regulation<br>Peroxisomal malate dehydrogenase; catalyzes interconversion of malate and oxaloacetate; involved in the glyoxylate cycle; mutation in human homolog MDH2 causes early-onset severe encephalopathy<br>Essential subunit of the Dam1 complex (aka DASH complex); cooperates with Dam1p and Pse1p to connect the DASH complex with microtubules (MT); couples kinetochores to the force produced by MT depolymerization thereby aiding in chromosome segregation; is transferred to the kinetochore prior to mitosis<br>Mitochondrial inner membrane protein; required for assembly of cytochrome c oxidase |
| *YOL095C* | *HMI1* | R, L | | |
| *YBL039W-B* | *MIN6* | R, M | | |
| *YBR018C* | *DYN2* | R, M | | |
| *YBR089C-A* | *NHP6B* | R, M | | |
| *YBR253W* | *SRB6* | R, M | | |
| *YDL078C* | *MDH3* | R, M | | |
| *YGL061C* | *DUO1* | R, M | | |
| *YJL003W* | *COX16* | R, M | | |
| *YML047C* | *PRM6* | R, M | | Potassium transporter that mediates K+ influx; activates high-affinity Ca2+ influx system (HACS) during mating pheromone response; expression up-regulated in response to alpha factor; regulated by Ste12p during mating; localized to sites of polarized growth; member of a fungal-specific gene family; PRM6 has a paralog, KCH1, that arose from the whole genome duplication |
| *YNL326C* | *PFA3* | R, M | | Palmitoyltransferase for Vac8p; required for vacuolar membrane fusion; contains an Asp-His-His-Cys-cysteine rich (DHHC-CRD) domain; autoacylates; required for vacuolar integrity under stress conditions<br>Putative protein of unknown function; expression is cell-cycle regulated, Azf1p-dependent, and heat-inducible; YNR014W has a paralog, YMR206W, that arose from the whole genome duplication |
| *YNR014W* | *YNR014W* | R, M | | |
| *YOL147C* | *PEX11* | R, M | | Peroxisomal protein required for medium-chain fatty acid oxidation; also required for peroxisome proliferation, possibly by inducing membrane curvature; localization regulated by phosphorylation; transcription regulated by Adr1p and Pip2p-Oaf1p<br>Putative protein of unknown function; green fluorescent protein (GFP)-fusion protein localizes to both the cytoplasm and the nucleus |
| *YEL025C* | *YEL025C* | S | | |
| *YFL030W* | *AGX1* | S | | Alanine:glyoxylate aminotransferase (AGT); catalyzes the synthesis of glycine from glyoxylate, which is one of three pathways for glycine biosynthesis in yeast; similar to mammalian and plant alanine:glyoxylate aminotransferases; human homolog AGXT complements yeast null mutant<br>Conserved zinc-finger domain protein involved in pre-mRNA splicing; critical for splicing of nearly all intron-containing genes; required for assembly of U4 snRNA into the U4/U6 particle |
| *YFR005C* | *SAD1* | S | | |
| *YGR215W* | *RSM27* | S | | Mitochondrial ribosomal protein of the small subunit |
| *YKL006W* | *RPL14A* | S | | |

| | | | | |
|---|---|---|---|---|
| *YLR185W* | *RPL37A* | S | | Ribosomal 60S subunit protein L14A; N-terminally acetylated; homologous to mammalian ribosomal protein L14, no bacterial homolog; RPL14A has a paralog, RPL14B, that arose from the whole genome duplication<br>Ribosomal 60S subunit protein L37A; required for processing of 27SB pre-rRNA and formation of stable 66S assembly intermediates; homologous to mammalian ribosomal protein L37, no bacterial homolog; RPL37A has a paralog, RPL37B, that arose from the whole genome duplication<br>Component of the exomer complex; the exomer which also contains Csh6p, Bch1p, Bch2p, and Bud7, is involved in the export of select proteins, such as chitin synthase Chs3p, from the Golgi to the plasma membrane; interacts selectively with the activated, GTP-bound form of Arf1p; Chs5p is the only protein with a BRCT domain that is not localized to the nucleus<br>Subunit of the RNA polymerase II mediator complex; associates with core polymerase subunits to form the RNA polymerase II holoenzyme; component of the Middle domain of mediator; required for regulation of RNA polymerase II activity; relocalizes to the cytosol in response to hypoxia<br>Transferrin receptor-like protein; functions with Tre1p to regulate ubiquitination and vacuolar degradation of the metal transporter Smf1p; inviability of null mutant in systematic studies is due to proximity to CDC31; TRE2 has a paralog, TRE1, that arose from the whole genome duplication |
| *YLR330W* | *CHS5* | S | | |
| *YNR010W* | *CSE2* | S | | |
| *YOR256C* | *TRE2* | S | | |
| *YPL158C* | *AIM44* | S | | Regulator of Cdc42p and Rho1p; regulates AMR closure through Hof1p; inhibits Cdc42-dependent Cla4 activation at the division site, to prevent budding in the old bud neck; recruits Nis1p and Nba1p to the division site with Nap1 and the Rax1p-Rax2p dependent inheritance of Nis1p and Nba1p to bud scars to prevent division site repolarization; keeps Rho1p at the division site after AMR contraction to control secondary septum formation; relocalizes from bud neck to cytoplasm upon replication stress |
